# The characteristics of intracellular Ca^2+^ signals *in vivo* necessitate a new model for salivary fluid secretion

**DOI:** 10.1101/2020.12.30.424864

**Authors:** Takahiro Takano, Amanda M. Wahl, Kai-Ting Huang, John Rugis, James Sneyd, David I. Yule

## Abstract

Salivary fluid secretion involves an intricate choreography to result in the trans-epithelial movement of NaCl and water into the acinus lumen. Current models are based on experimental observations in enzymatically isolated cells where the Ca^2+^ signal invariably propagates globally and thus appears ideally suited to activate spatially separated Cl and K channels. We monitored Ca^2+^ signals and salivary secretion in live mice expressing GCamp6F, following stimulation of the nerves innervating the submandibular gland. Consistent with *in vitro* studies, Ca^2+^ signals were initiated in the apical endoplasmic reticulum. In marked contrast to *in vitro* data, highly localized trains of Ca^2+^ transients that failed to propagate from the apical region were observed. Following stimuli optimum for secretion, large apical-basal gradients were elicited. Given this incompatibility to the previous model, a new mathematical model was constructed to explain how salivary secretion can be efficiently stimulated by apically localized Ca^2+^ signals.

## Introduction

Salivary fluid secretion is vital to the health of the oral cavity [37]. This is most clearly appreciated in disease states which result in hypofunction where patients experience difficulty chewing and swallowing food, speaking and are susceptible to oral infections that result in dental caries [29]. Fluid secretion from the salivary gland requires the trans epithelial movement of Cl^-^ across polarized acinar cells [16, 30, 31, 49]. The current, widely accepted model for fluid secretion posits that Cl^-^ are transported against their electrochemical gradient by the Na^+^/K^+^/2Cl^-^ cotransporter, NKCC1 at the basolateral plasma membrane (PM) [15] and subsequently enters the lumen of the acinus across the apical PM through TMEM16a chloride channels [39]. Cl^-^ accumulation imparts a negative potential to the lumen relative to the basolateral PM, which draws Na^+^ paracellularly through tight junctions [31]. Subsequently, water follows osmotically to generate the primary saliva which is modified by the ductal epithelia. An elevation in cytosolic [Ca^2+^] following neurotransmitter release plays a central role in the stimulation of fluid secretion from salivary gland acinar cells [3, 4, 18, 38, 53]. Acetylcholine (ACh) is the primary neurotransmitter released following parasympathetic input to the major salivary glands after gustatory and olfactory stimulation [28]. ACh results in activation of muscarinic receptors, G-protein coupled stimulation of inositol 1,4,5 trisphosphate (IP_3_) production and subsequent Ca^2+^ release from endoplasmic reticulum stores via IP_3_ receptors (IP_3_R) [17, 24, 34]. The ER stores are subsequently refilled by store operated Ca^2+^ entry across the plasma membrane (PM), which is required for sustained salivary secretion [14, 44]. To stimulate saliva flow, the primary effector of the increase in [Ca^2+^] are apically localized TMEM16a channels [5, 6, 10, 39]. As Cl^-^ moves across the apical PM, the membrane potential approaches the equilibrium potential for Cl^-^, however, Ca^2+^ also increases the activity of K channels localized predominately at the basolateral PM to maintain the membrane potential to favor continued Cl^-^ exit and secretion [27, 32, 33, 40]. Over the past several decades, the spatiotemporal properties of the agonist-stimulated Ca^2+^ signal have been described in preparations of enzymatically isolated preparations of acinar cells or lobules [36, 52] loaded with fluorescent Ca^2+^ indicators and imaged by time resolved wide field or confocal microscopy. In total, these studies indicate that the agonist stimulated Ca^2+^ signals exhibit characteristics which appear ideally tuned to activate these spatially separated Ca^2+^ dependent ion channels required for secretion. For example, stimulation with muscarinic agonists, or more directly by photolysis of caged-IP_3_, invariably results in the Ca^2+^ signal initiating in the extreme apical ER followed by a rapid Ca^2+^ wave which results in globalization of the signal throughout the cytoplasm [11, 12, 19, 22, 48, 54]. The initiation sites, termed the “trigger zone”, correspond to the localization of the majority of type-2/3 IP_3_R [24, 34], in ER in close apposition to TMEM16a in the PM. The global spread is consistent with Ca^2+^-induced Ca^2+^ release through IP_3_R and ryanodine receptors (RyR) [54] to allow the Ca^2+^ signal to efficiently activate the majority of K channels present on the basolateral PM. At low agonist concentrations, the cycle of apical Ca^2+^ release, global spread and subsequent Ca^2+^ sequestration is repeated, such that temporally, the Ca^2+^ signal is manifested as a series of oscillations with a relatively slow period (5-8/minute) [12]. At higher concentrations of agonist, the initial globalization is maintained to produce a “peak and plateau” type temporal response.

While *in vitro* studies of Ca^2+^ signaling have provided a wealth of information to greatly inform our models of salivary secretion, there are general caveats which apply to these data. Primarily, the cells are studied out of a physiological context with no extracellular matrix, basement membrane and with probable disruption of cellular junctions as a result of the enzymatic isolation. A further complication is that isolated cells are typically exposed for a period of time to a constant concentration of agonist which in many cases is a non-hydrolysable analog, as opposed to the neural delivery of transmitter occurring in a pulsatile fashion that is subject *in vivo* to enzymatic degradation. Whether these factors influence the spatiotemporal characteristics of Ca^2+^ signals in salivary glands is unknown. Additionally, since only indirect measurements of fluid secretion inferred from volume changes can be made in isolated acinar cell preparations [45], it is unclear how the concentration-dependent characteristics of stimulated Ca^2+^ signals *in vitro* relate to the physiological stimulation of fluid secretion *in vivo*.

In this study we generated mice expressing a genetically encoded Ca^2+^ indicator specifically in exocrine acinar cells and imaged the Ca^2+^ signals in submandibular glands in living mice by intravital multiphoton (MP) microscopy. Notably, stimulation of the endogenous neural input to the gland induces Ca^2+^ signals with major spatiotemporal properties which are distinct from those observed in isolated acinar cells. Specifically, following moderate stimulation, apically localized Ca^2+^ signals are generated which did not propagate globally. Furthermore, the apical signals consist of rapid oscillations. At stimulation intensities optimum for secretion, a significant apical-basal standing Ca^2+^ gradient was also established. These striking disparities necessitate a modification of our previous model [34, 42] to incorporate the physiological characteristics of Ca^2+^ signals *in vivo* and thus provide a revised framework to better understand salivary fluid secretion.

## Results

To generate mice expressing a fluorescent Ca^2+^ indicator in exocrine acinar cells, homozygous mice expressing the fast genetically encoded Ca^2+^ indicator GCaMP6f [13] floxed by a STOP cassette, were crossed with heterozygous tamoxifen-inducible Mist1 Cre mice [26]. The STOP codon was subsequently excised following tamoxifen gavage (Fig. 1A). The expression of GCaMP6f was investigated *in vivo* in anaesthetized mice following surgery to expose the submandibular salivary glands (SMG). One SMG was lifted and placed on a small platform with a coverslip placed on top of the gland (Fig 1B). Bipolar stimulation electrodes were inserted to the duct bundle connecting to the imaged SMG tissue, in order to stimulate nerves innervating the SMG. A cover glass was placed on top of the SMG to gently hold the tissue and to maintain a saline solution bathing the gland. MP imaging in serial z sections through the gland revealed expression of GCaMP6f exclusively in acinar cells. GCaMP6f expression was uniformly observed in essentially all acinar cells while GCaMP6f expression was not evident in any other cell type such as blood vessels, nerves, myoepithelial cells or ductal cells. Voids with lack of expression of GCaMP6f were evident in z planes distal from the surface of the gland likely representing the localization of ductal structures (Fig 1C).

**Fig. 1.**
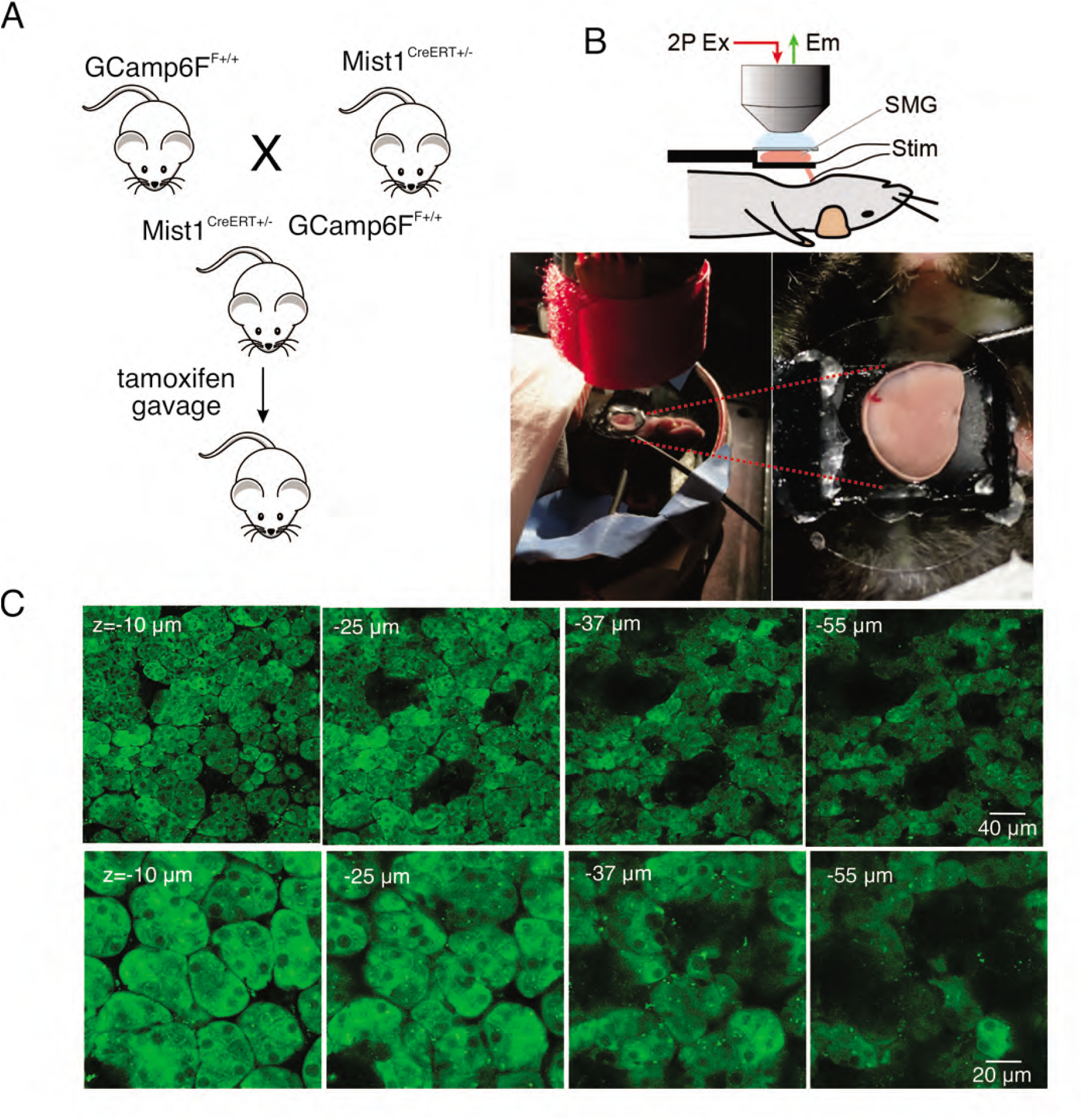
*In vivo* SMG Ca^2+^ imaging. **A**. Generation of transgenic mice expressing the GCaMP6f gene driven by Mist1 promotor. GCaMP6f is selectively expressed in acinar cells. **B**. Animal preparation for *in vivo* imaging. A SMG was lifted and placed on a small platform, then held with a coverslip on top. Stimulation electrodes were inserted to a duct bundle that connect SMG to the body to stimulate nerves that innervate to SMG. **C**. A series of z-projection images of SMG with GCaMP (green) expressed in acinar cells.

We next monitored changes in GCaMP6f fluorescence by intravital microscopy in live animals using MP excitation. Images were acquired at 10 frames per second as described in Methods (each image a 3 frame average). Under basal conditions, prior to stimulation, the cells were quiescent (Fig. 2B). Electrical stimulation at various stimulus strengths delivered to nerves innervating the SMG (1-100 Hz, 12 s initiated at 10s) induced increases in intracellular [Ca^2+^] in acinar cells (Fig. 2A,B). Following 1 Hz stimulation, a minority of the cells exhibited small, rapid changes in fluorescence, (n = 4 fields from 4 animals, Fig. 2A,B), while most cells were refractory. Following higher frequency stimulation (3-10 Hz), the majority of the cells in the field responded and the amplitude of the [Ca^2+^] signal increased as the stimulation strength was increased (Fig. 2A,B). During the 12 s of stimulation, overall fluorescence gradually increased and immediately returned to baseline after the termination of the stimulation (Fig. 2A,B). The maximum [Ca^2+^] signal evoked during the 12 s stimulation progressively increased and was maintained from 1 to 10 Hz throughout the period of stimulation, but this trend did not hold following higher frequency stimulation (n = 4 fields from 4 animals, Fig. 2C). Following stimulation at 30-100 Hz, the [Ca^2+^] increase was not maintained for the duration of stimulation. Under these conditions, the [Ca^2+^] rapidly peaked in the initial 1-3 s then decreased thereafter (Fig. 2A,B), probably as a result of exceeding the maximal firing rate of the neurons innervating the SMG. Of note, the stimulation likely led to increased neural activity specifically, rather than stimulating acinar cells directly, as a consequence of current leak through the saline depolarizing nerve endings, as the contralateral SMG were unresponsive to even the highest 100 Hz nerve bundle stimulation of the ipsilateral gland (p > 0.1 compared to without stimulation, n=4 fields from 4 animals, Fig. 2C).

**Fig. 2.**
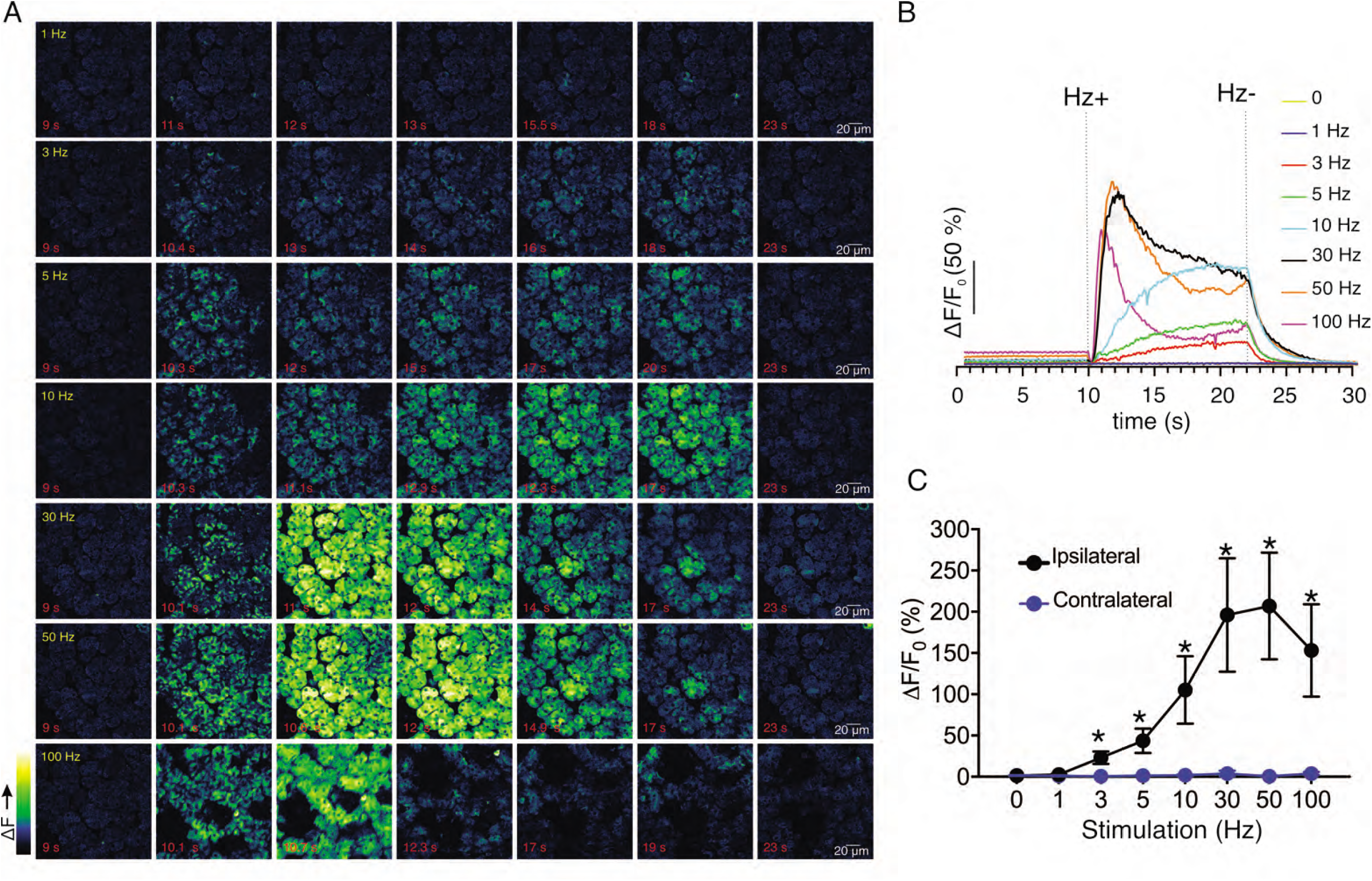
Ca^2+^ signals evoked in response to nerve stimulation. **A**. Time-course images of Ca^2+^ (baseline fluorescence was subtracted) in response to 1-100 Hz stimulations. **B**. Mean response of the entire field of view to stimulation at the indicated frequency. N = 4 fields from, 4 animals. **C**. A summary of peak Ca^2+^ increases during each stimulation. Stimulation was only to the ipsilateral SMG, which contralateral SMG failed to respond. N = 4 fields from 4 animals. Mean ± sem. *p <0.05 *vs*. contralateral gland, t test.

To further confirm the physiological consequence of nerve stimulation, we monitored saliva secretion under identical stimulus conditions. There was no significant secretion activated by 1-3 Hz nerve stimulation for 1 min (p > 0.05 compared to the absence of nerve stimulation, n = 4 animals), but saliva secretion progressively increased following 5-10 Hz nerve stimulation (p < 0.05 compared to the absence of stimulation, n = 4 animals) (Fig. 3A). Higher frequency stimulation did not result in augmented secretion, likely reflecting failure to sustain the rise in [Ca^2+^] (Fig. 2A). At moderate stimulation intensity (0-10 Hz), there was a linear correlation between the peak [Ca^2+^] increases by stimulation and saliva secretion (r^2^ = 0.995, n = 4 animals, Fig. 3B). These stimulus paradigms (1-10 Hz), therefore likely represent the range of physiological stimulation of both evoked Ca^2+^ signals and the resulting saliva secretion and thus characterization of these signals, represent the focus of the remainder of the study.

**Fig. 3.**
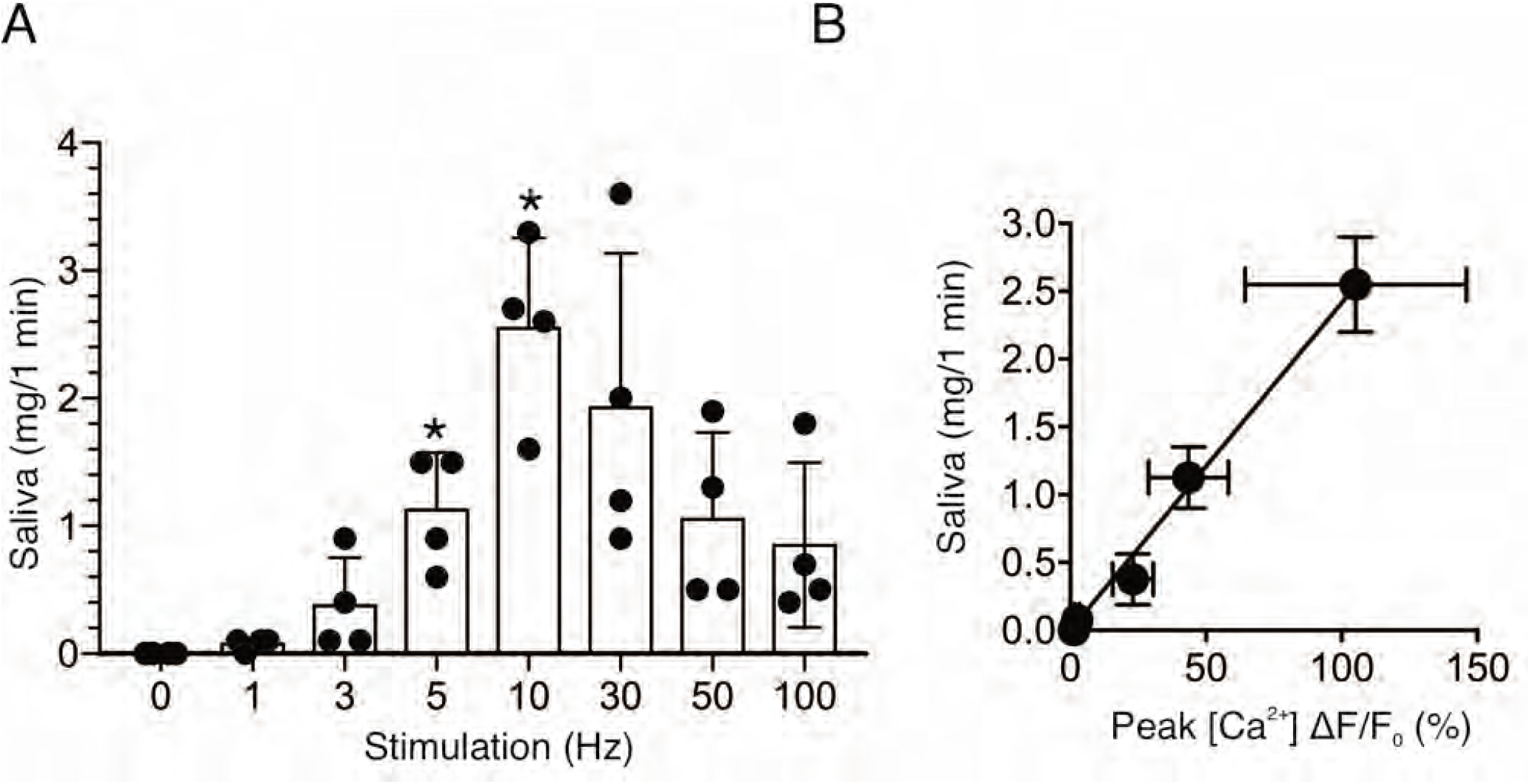
Saliva secretion following nerve stimulation. **A**. A summary histogram of total saliva secretion following 1-min stimulations at the indicated frequency. N = 4 animals. Mean ± sem. **B**. A correlation plot of peak Ca^2+^ (shown in Fig. 2C) vs. saliva secretion, which showed a linear regression of R^2^ = 0.995 (black line). *p <0.05 *vs*. no stimulation ANOVA with Dunnett test.

Following nerve stimulation at low stimulation strengths (1-5 Hz), we observed both the recruitment of responding cells and an increase in the amplitude of the [Ca^2+^] response (Fig. 4). To determine more closely how the increased stimulation intensity impacts the characteristics of Ca^2+^ responses to result in increased saliva secretion, a software-based work-flow was designed to post-process the image series. Imaged fields were sub-divided into 8 by 8 equal quadrants to yield 64 grid squares (Fig. 4C,D). Each grid was 32 x 32 µm, and typically included a portion of an acinus consisting of 1-5 cells and was used to approximate the spatial and temporal behavior of individual cells or lobules in the field. In order to analyze the activity in each region, the image was then processed to show percent ΔF/F for each time-lapse frame in the sub-divided regions (Fig. 4D,E). Using this general scheme, we compared information obtained from 8 x 8 grids, 16 × 16 grids and randomly manually selected regions of interest representing either acinar clusters or single acinar cells. Each scheme produced essentially equivalent data, ultimately validating the use of the 8 x 8 grid (Supplemental Fig. 1). At 1 Hz stimulation, the activity was sporadic, with the responding regions tending to give multiple responses while non-responding regions remained quiescent throughout the stimulus period (Fig. 4F). Overall, 34.0 ± 5.1% (n = 19 fields from 8 animals) of the sub-regions showed significant responses during the stimulation (Fig. 4G). Moreover, the increase in [Ca^2+^] in the responding regions exhibited a prolonged latency prior to the initial rise following stimulation (6.72 ± 0.79 s, n = 19, Fig. 4H). Stimulation at higher frequencies progressively recruited more grid regions and shortened the latency to the initial response (Fig. 4G, H). At 10 Hz, virtually all regions responded to the stimulation (98.5 ± 0.6%, n = 19) with minimal latency (1.70 ± 0.27 s, n = 19) (Fig. 4G, H). These data suggest that the increase in overall [Ca^2+^] and secretion by rising stimulation strength was at least in part due to the recruitment of responding cells and a faster time for cells to respond. Further, the amplitude of the maximal increase in [Ca^2+^] in the subdivided grids also increased as the strength of the stimulation was augmented (Fig. 4I), indicating that enhancement of individual cellular [Ca^2+^] responses also contribute to the overall increase of [Ca^2+^] amplitudes.

**Fig. 4.**
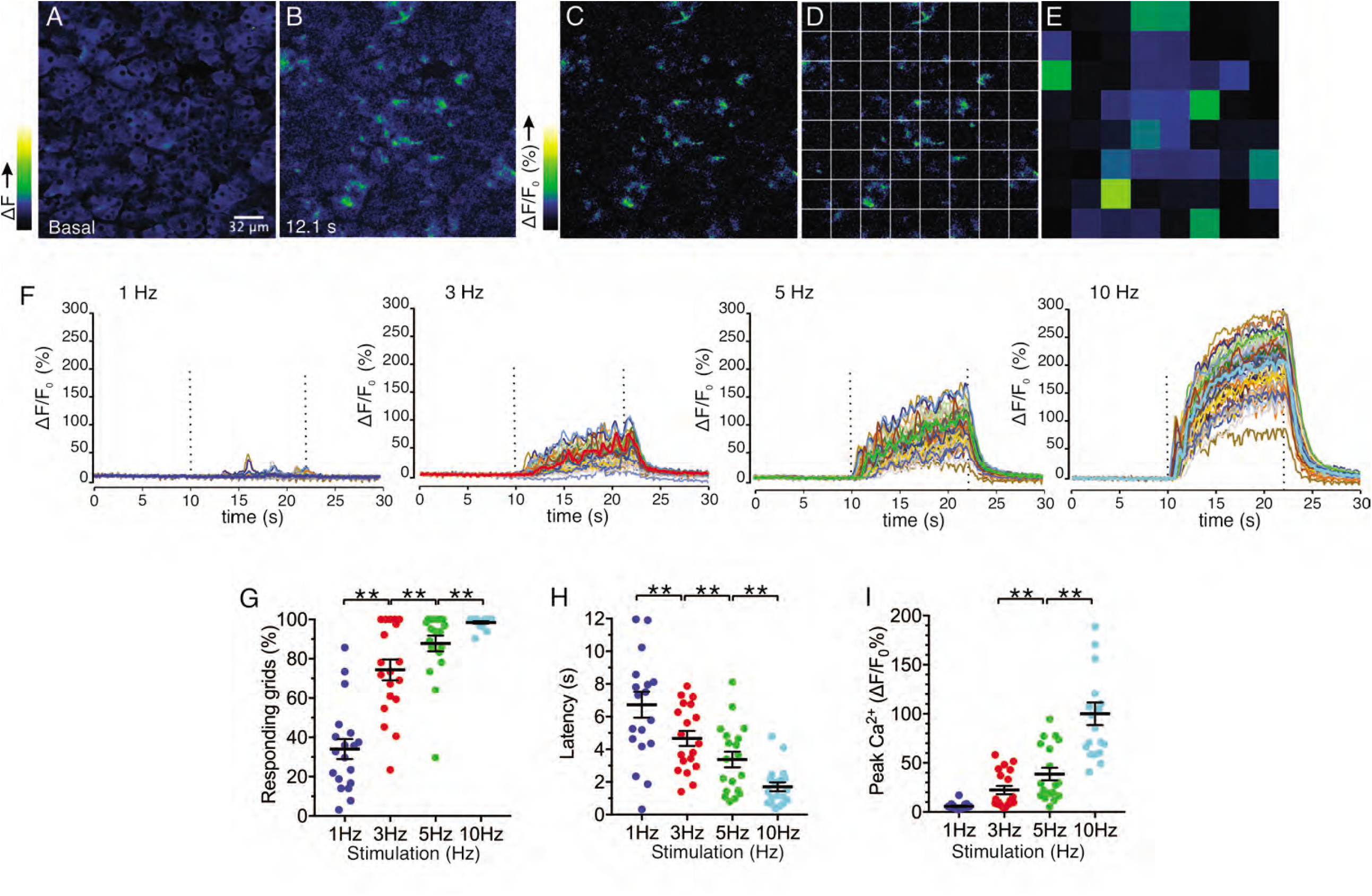
Spatial analysis of Ca^2+^ responses to various stimulations. **A-B**. An imaging field before (**A**) and during (**B**) a stimulation. **C-D**. An imaging field was divided to grids of 32 x 32 µm, to yield 64 regions. **E**. The average intensity per grid was obtained in each time frame. **F**. Representative time-course plots of all 64 grids before, during, and after 1-10 Hz stimulations. Thick colored line represents the mean of the 64 grids for each stimulation. **G**. A summary plot of average percent of responding grids in a field by the stimulations. N = 19 from 8 animals. Mean ± sem. **H**. A summary plot of average latencies to initiation of Ca^2+^ increases in all grids. Non-responding grids were excluded for the data. Mean ± sem. N = 19 from 8 animals. **I**. A summary plot of average peak Ca^2+^ increases in all grids. Mean ± sem. N = 19 from 8 animals. **P<0.01 ANOVA with Tukey test.

We observed that increased stimulus strength resulted in both a higher number of responding regions and a higher peak [Ca^2+^] amplitude in individual grids. We next examined whether grids which responded at lower stimulus intensity also strongly responded to higher frequency stimulation. The alternative being that the spatial pattern of responding grid regions differed with increasing strengths of stimulus. Grids in an imaging field were ranked based on the peak [Ca^2+^] evoked by 5 Hz stimulation and then re-sorted based on progressively increasing amplitude (Fig. 5A-B). We subsequently sorted the data obtained from 1, 3, and 10 Hz stimulation from the identical field in the grid rank order established for 5 Hz stimulation (Fig. 5C). This analysis demonstrated a clear pattern whereby, at each stimulus strength, particular regions of the field of view were most susceptible to stimulation, such that grids giving the largest amplitude response at 1 Hz similarly responded by displaying the largest amplitude signals at 3, 5 and 10 Hz stimulation. (n= 5 fields from 4 animals) (Fig. 5D).

**Fig. 5.**
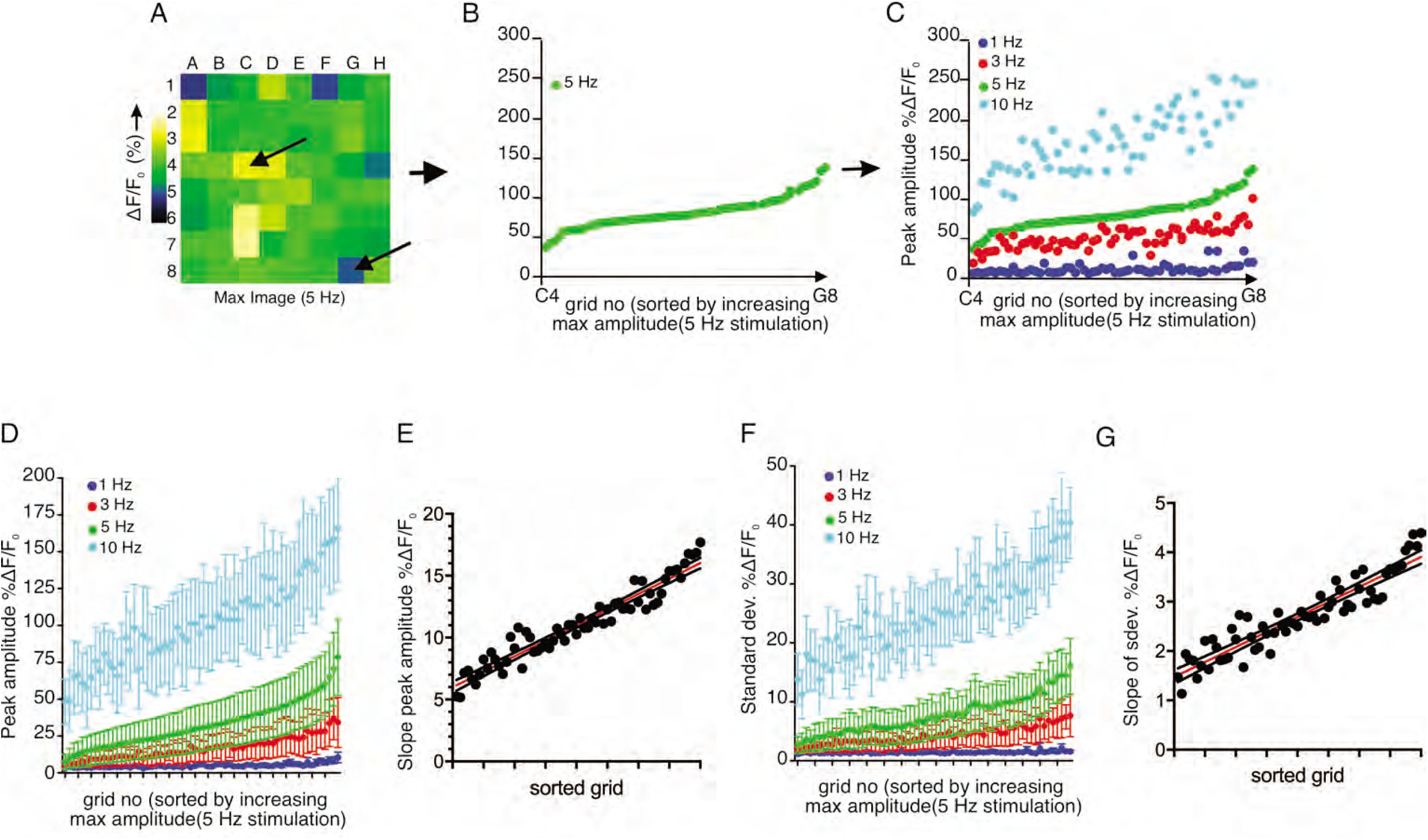
Each sub-divided field possesses unique sensitivity. **A**. An example image of a processed imaging field showing peak intensities during a stimulation in each grid. The grids consist of lowest intensity (arrow, grid A1) to highest (arrow, grid C7). **B**. A representative plot of peak Ca^2+^ increases for each grid in a field to a 5 Hz stimulation. The grids in x axis were reordered from the lowest intensity to the left to the highest on the right. **C**. Different stimulation strengths were applied to the same field, and the responses were plotted according to the order indexed by the responses to 5 Hz. **D**. A summary plot of peak Ca^2+^ increases in each grid sorted by responses to 5 Hz. Mean ± sem. N = 5 from 4 animals. **E**. A correlation plot of sorted grid, from low peak Ca^2+^ to high, vs. degree of Ca^2+^ increases in response to higher stimulation. Linear regression with R^2^= 0.920. N = 5 from 4 animals. **F**. A summary plot of standard deviations of Ca^2+^ changes during stimulations in each grid sorted by responses to 5 Hz. N = 5 from 4 animals. Mean ± sem. **G**. A correlation plot of sorted grids, from low peak Ca^2+^ to high, vs. degree of standard deviation of Ca^2+^ flux in response to higher stimulation. Linear regression with R^2^ = 0.866. N = 5 from 4 animals.

The extent of the increase in [Ca^2+^] following greater strengths of stimulation was not uniform throughout individual grids. High-responding grid regions showed linearly enhanced increases following stimulation (R^2^ = 0.9197, n = 4 fields from 4 animals), while the increase in Ca^2+^ signal following increased stimulation in less responsive regions was proportionately smaller (Fig. 5E). This trend was not limited to the peak amplitude attained. During the 12 s stimulation, grid regions exhibited [Ca^2+^] fluctuations superimposed upon an elevated plateau (Fig. 4F). We evaluated the extent of this fluctuation by comparing the standard deviation of the ΔF/F signal occurring during the stimulation period. With this analysis, a higher standard deviation reflects greater variability in the stimulated change in [Ca^2+^]. When the 5 Hz data was sorted according to the peak [Ca^2+^] ranking as in Fig. 5D, the variation of grids with 5 Hz stimulation also demonstrated that low-responding grids had smaller variability than high-responding grids (n = 4 fields from 4 animals) (Fig. 5F). Likewise, the same trend was exhibited at each stimulation strength, with a linearly increasing rate of enhancement as intensity increased (R^2^ = 0.8664, n = 4) (Fig. 5G). Therefore, an increased fluctuation in the response is also a property of sensitive, high responding regions.

Stimulation of salivary fluid secretion is predominately mediated through parasympathetic input as a result of the action of ACh released from nerve endings acting on muscarinic M3 receptors on acinar cells. To gain direct evidence that neural release of ACh was responsible for the changes in [Ca^2+^] observed, we systemically administered a cholinesterase inhibitor to block the clearance of ACh and effectively increase the local [ACh]. Physostigmine (0.1 mg/kg body weight *IP*) enhanced the peak amplitude of the Ca^2+^ signals following stimulation (Fig. 6A-D), indicating that acinar cell [Ca^2+^] changes reflect the extracellular availability of acetylcholine released by stimulation. The augmentation in the Ca^2+^ signal was not statistically significant at 1 Hz stimulation (p > 0.05, n = 5 fields from 4 animals), but became more prominent at 3 Hz with shorter latency, an increase in the number of responding grids and peak amplitude (p < 0.05 in % response and peak ΔF/F, n = 5) and following 5 Hz stimulation (p < 0.05 in latency and peak ΔF/F, n =5 fields from 4 animals) (Fig. 6E-G). At 10 Hz stimulation, no further increase in the grid responsiveness or latency was observed, reflecting the fact that all grids were active and exhibited minimal latency at this stimulus strength, however the peak ΔF/F showed a robust enhancement (p < 0.01, n = 5) (Fig. 6E-G). While these data do not rule out the involvement of additional neurotransmitters, nevertheless they confirm a prominent role for ACh in mediating the increase in [Ca^2+^] observed following neural stimulation.

**Fig. 6.**
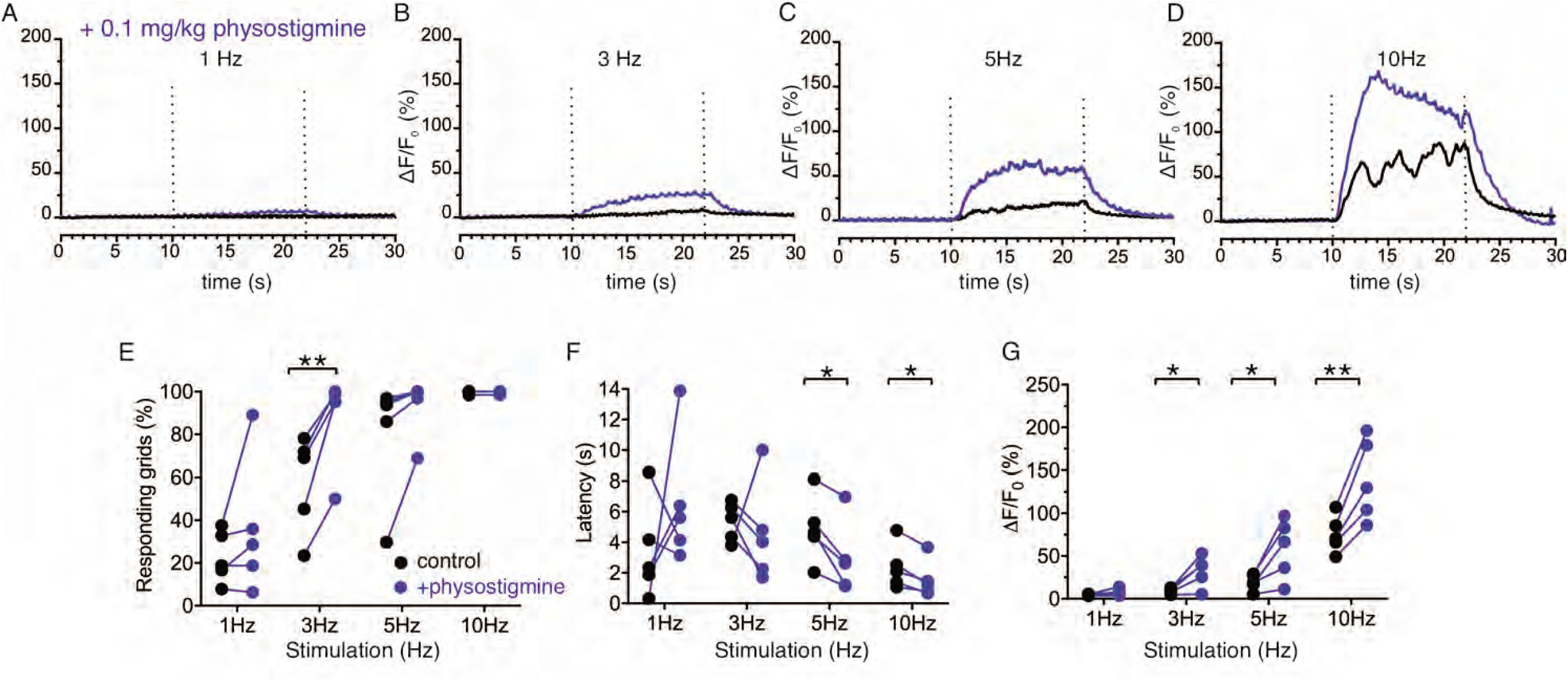
Enhancement of Ca^2+^ responses by cholinesterase inhibition. **A-D**. Representative average traces from 64 grids of changes in Ca^2+^ responses in a same field before (black) and after (blue) physostigmine. **E**. A summary plot of average percent of responding grids in a field by the stimulations in paired fields. N = 5 from 4 animals. Mean ± sem. **F**. A summary plot of average latency to initiation of Ca^2+^ increases in all grids in paired experiments. Non-responding grids were excluded for the data. N = 5 from 4 animals. Mean ± sem. **G**. A summary plot in paired experiments of average peak Ca^2+^ rise in all grids. N = 5 from 4 animals. Mean ± sem. *P<0.05; **P<0.01 Paired t test.

While the previous grid-based analysis yielded important information relating to the general properties of the evoked signals within an imaged field, this approach was not suited to provide spatial information on a sub-acinus or lobule scale. To interrogate the sub-cellular spatial characteristics of the Ca^2+^ signals evoked by neural stimulation, we generated standard deviation (SD) images from the stimulated portion of the image series. This analysis revealed that at all physiological stimulus strengths the increase in [Ca^2+^]i was highly heterogeneous, with the largest increases in signal occurring in the apical aspects of acinar clusters (Supp. Fig 2). Figure 7A illustrates a field in which an acinar cluster is highlighted prior to neural stimulation at 5 Hz (red box and image series in Fig 7B). In marked contrast to *in vitro* experiments, analysis of the image series demonstrated that the signal consisted of brief (< 2 sec in duration), repetitive increases in [Ca^2+^] (Fig 7B). The signals were highly localized, being initiated in, and almost entirely confined to the apical portions of individual cells (fig 7B-E) with only very rare, small elevations in the basal aspects of the cell. This localization is illustrated by SD images of the whole field (Fig 7D) and a topographical SD representation of the highlighted cluster (Fig 7D) together with analysis of the fluorescent changes in time from regions of interest manually assigned to an apical Region of Interest (ROI) (red box) and a basal ROI (black box) (Fig 7E). Consistent with tight coupling between acinar cells by gap junctions, signals often appeared to invade the apical region of neighboring cells (see images time stamped between 30.8-31.8 s in Fig 7B).

**Fig. 7.**
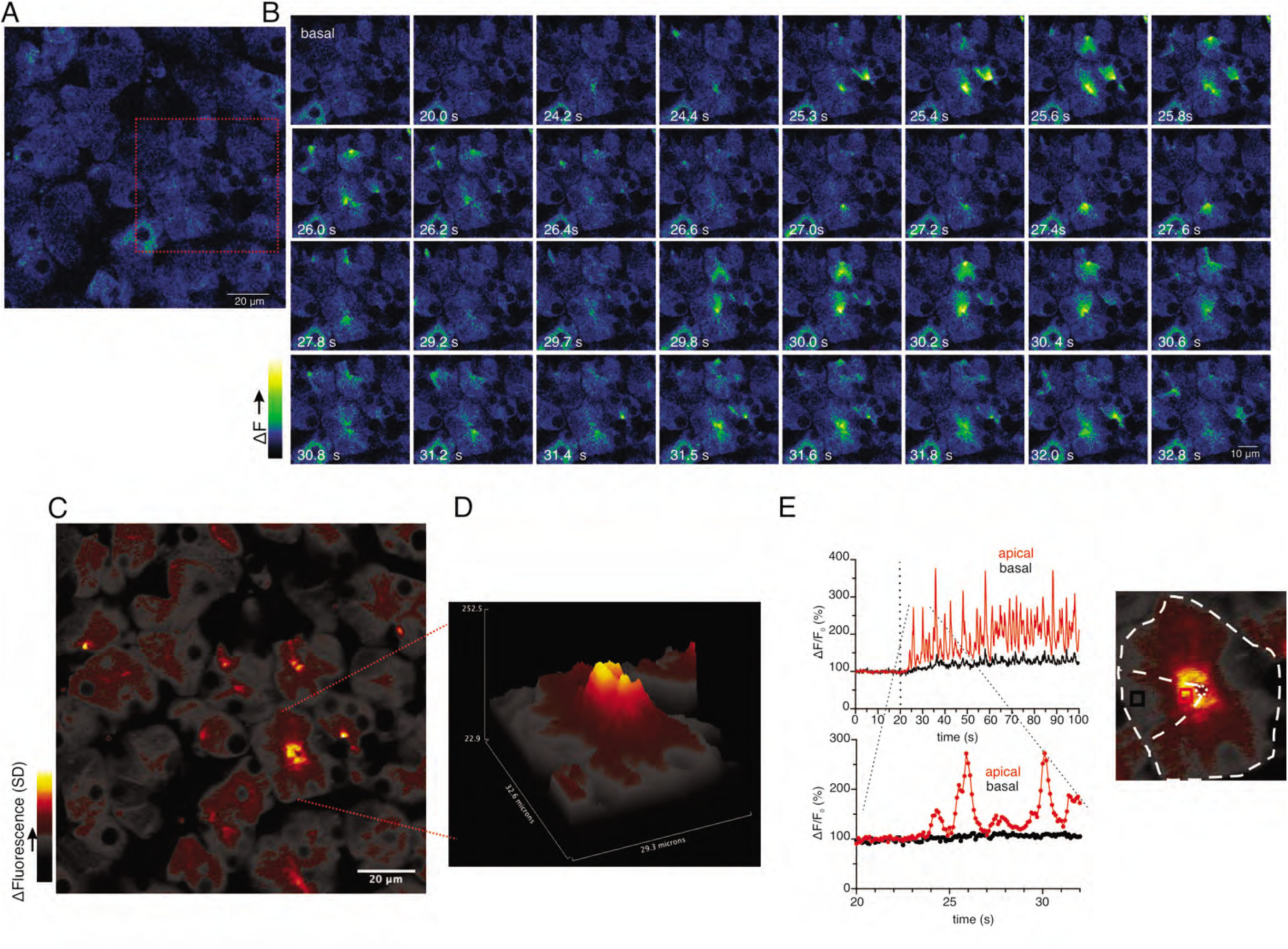
Ca^2+^ signals are restricted to the apical region during moderate stimulation. **A**. A representative imaging field prior to stimulation. **B**. Time-course images of a field containing an acinar cluster (red box in panel A) showing active Ca^2+^ signaling in the extreme apical portion in the cells. **C**. A processed image mapping standard deviation values during stimulation period in the field shown in panel A. **D**. A 3D plot of the SD image of the acinar cluster in B, showing large fluctuations in [Ca^2+^] only in the apical domain with minimal Ca^2+^ signal transmission to the basal aspects of the cell. **E**. Plot of Ca^2+^ changes in a single cell from the apical domain (red box in the image at right) and basal (black box in the image at right).

At 10 Hz stimulation, a rise in [Ca^2+^] was also rapidly initiated in the extreme apical region of the acinus (imaging field in Fig 8A with red box showing the highlighted cluster depicted in Fig 8B). In contrast to lower stimulus strengths, the evoked Ca^2+^ signals propagated to the basal aspects of the cell (Fig 8B-E). Notably, however the magnitude of the signal was invariably larger in the apical domain, approaching saturation of the indicator, throughout the entire duration of the stimulation. Effectively a standing apical-basal [Ca^2+^] gradient was established, as shown in the SD image of the field and the topographical representation of the SD of the highlighted region (Fig 8C/D), together with the analysis of manually positioned apical and basal ROIs (Fig 8E). In total, these data demonstrate that in contrast to experiments performed in *in vitro* preparations, physiological stimulation results in prominent, apical Ca^2+^ signals which do not invariably propagate and globalize to result in a homogeneous [Ca^2+^] throughout the acinar cell.

**Fig. 8.**
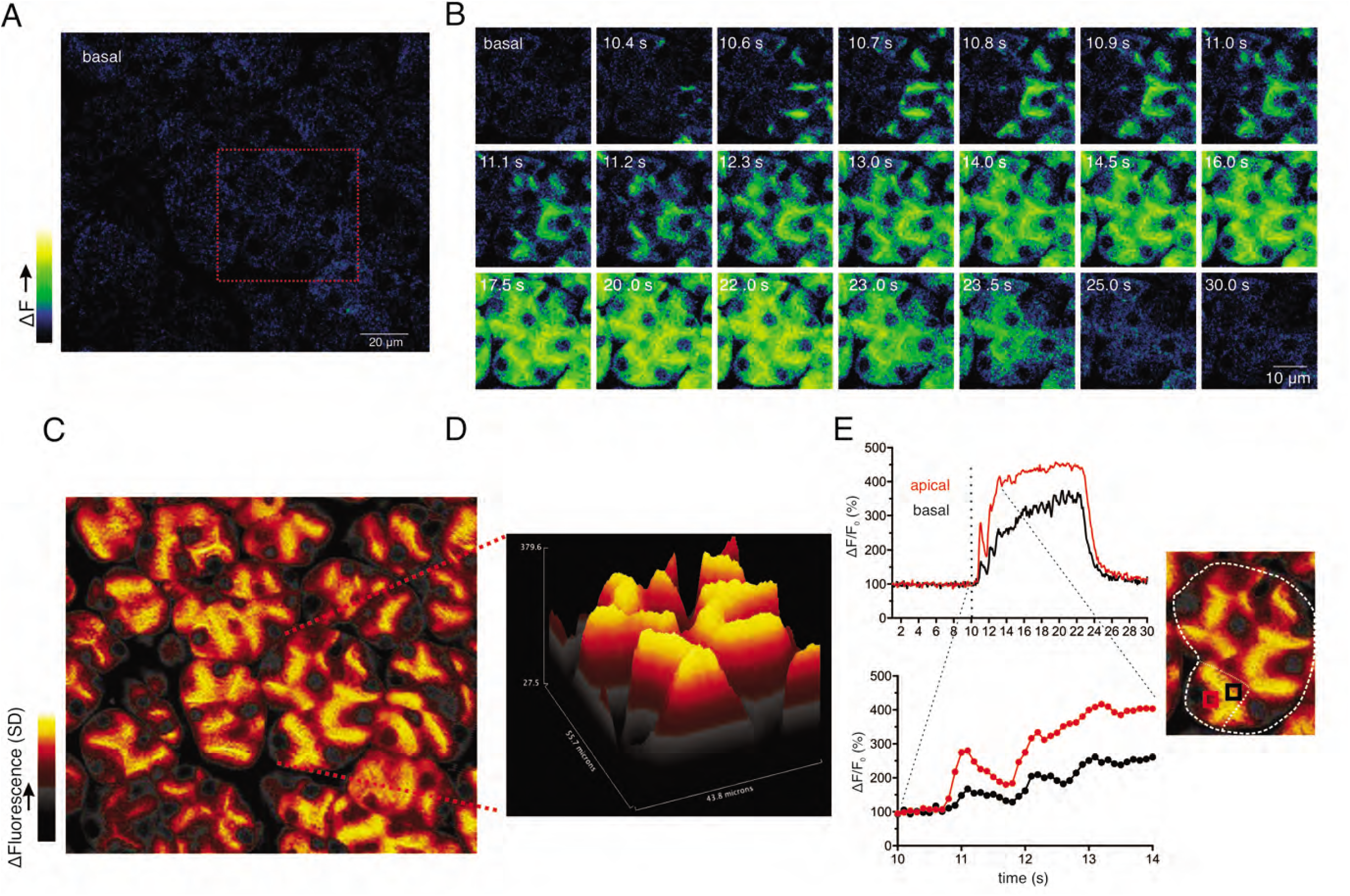
Apical to basal [Ca^2+^] gradients are established during optimal stimulation. **A**. A representative imaging field prior to stimulation. **B**. Time-course images of a field containing an acinar cluster (red box in panel A) depicting predominant Ca^2+^ changes in the apical portion of cells with smaller changes in the basal regions. **C**. A processed image mapping standard deviation values during the stimulation period in the whole field shown in panel A. **D**. A 3D plot of the SD image for the acinar cluster shown in B **E**. Tracings of Ca^2+^ changes in the apical domain (red box in the image at right) and basal aspects of a single cell from this cluster (black box in the image at right).

To further analyze the neurally evoked apical Ca^2+^ signals, Python software scripts were written in the Jupyter lab notebook environment to automate the designation of apical ROIs from masks generated from average or SD images. These apical ROIs were then subsequently used to quantitate the magnitude and frequency of signals within these regions (see Methods for details). Figure 9A shows an imaging field depicting the stimulated-basal average image for the field generated during the 12s of stimulation at 5 Hz from which 79 apical ROIs were generated from masks shown in Figure 9B/C. Figure 9D shows representative responses from 4 of the 79 ROIs to stimulation over the range of physiological stimulus strengths. Consistent with the previous grid analysis, few apical regions responded to 1 Hz stimulation while the majority of ROIs responded to stimulation at 5-10 Hz (see supplementary Movies 1-4). At both 3 and 5 Hz stimulation, individual apical ROIs typically responded by generating a rapid train of oscillations. A peak counting algorithm was used to analyze the frequency of the oscillations for each ROI. This demonstrated that the frequency of oscillations was markedly faster (∼1 Hz) when measured *in vivo* than previously described in isolated cells (∼0.15 Hz, this study, Fig 10) and further modestly increased between 3 and 5 Hz stimulation (Fig 9E; 79 ROIs experiment shown in Fig. 9A-C and pooled data from 13 individual fields from 5 animals (Fig 9F)). In contrast to the robust correlation between saliva secretion and the magnitude of the evoked Ca^2+^ signal, the relationship between secretion and the frequency of Ca^2+^ oscillation was less strong (R^2^ =0.824, Fig 9G).

**Fig. 9.**
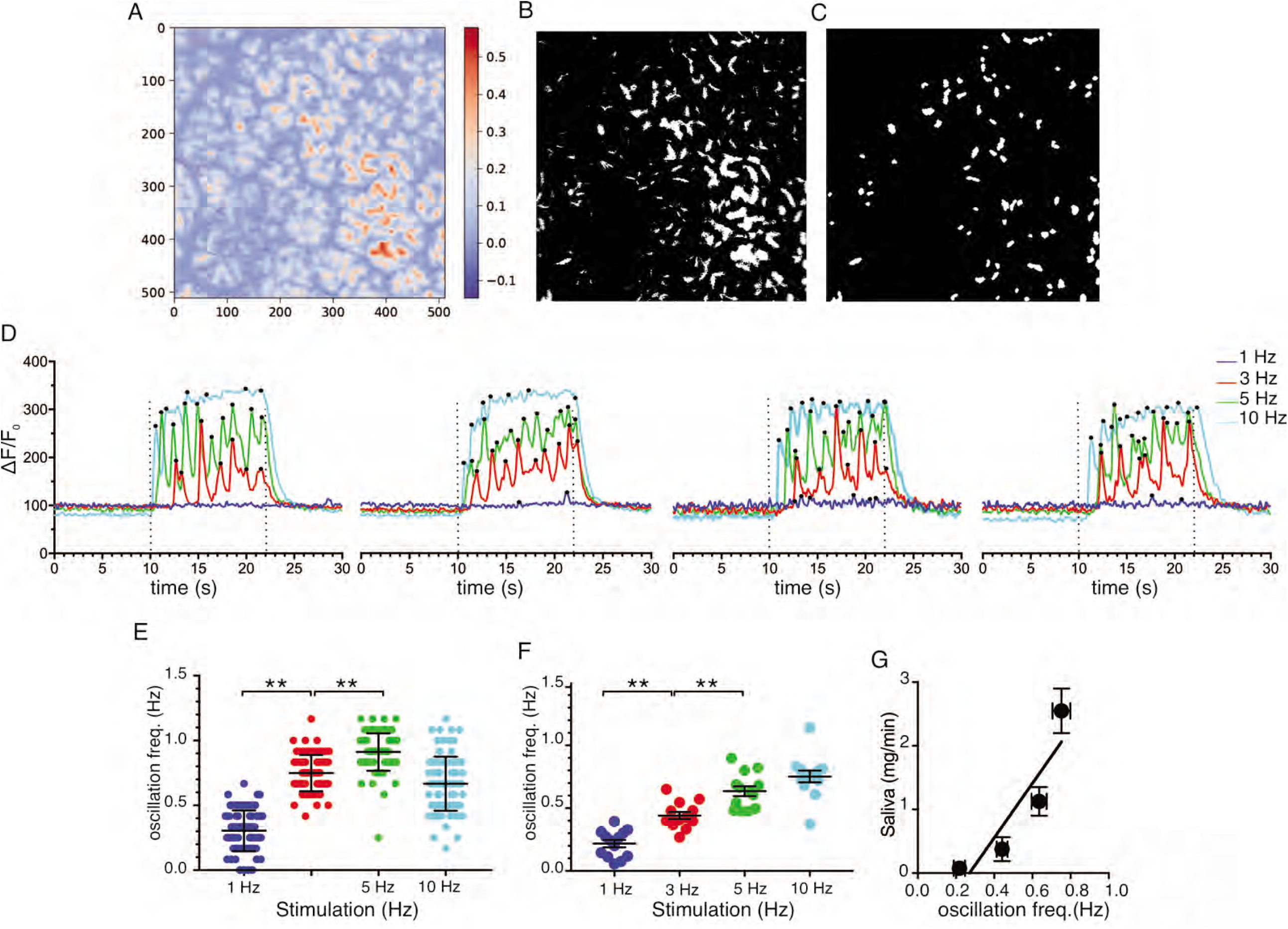
Ca^2+^ Oscillation frequencies mildly correlates with stimulation strengths. **A**. A representative image generated from averaging signal intensities during a 5 Hz stimulation, which highlights large fluorescence changes in the apical aspects of the cells. **B-C**. Mask generated from the image in panel A using Python scripts running in the Jupyter lab environment, as described in Methods. **D**. Representative Ca^2+^ responses from 4 of the ROIs defined in the image in panel C following stimulation at 1-10 Hz. Black dots designate positions of oscillation peaks detected by automated peak detector software written in python. **E**. A summary of oscillation frequency of all 79 ROIs from the image in panel A-D. Mean ± sem. **F**. A summary plot of oscillation frequencies in response to 1-10 Hz stimulation. N = 13 fields from 5 animals. Mean ± sem. **F**. A correlation plot of oscillation frequency vs. saliva secretion (shown in Fig. 3A), which showed a linear regression with R^2^ = 0.824 (black line). ** P<0.01 ANOVA with Tukey test.

**Fig. 10.**
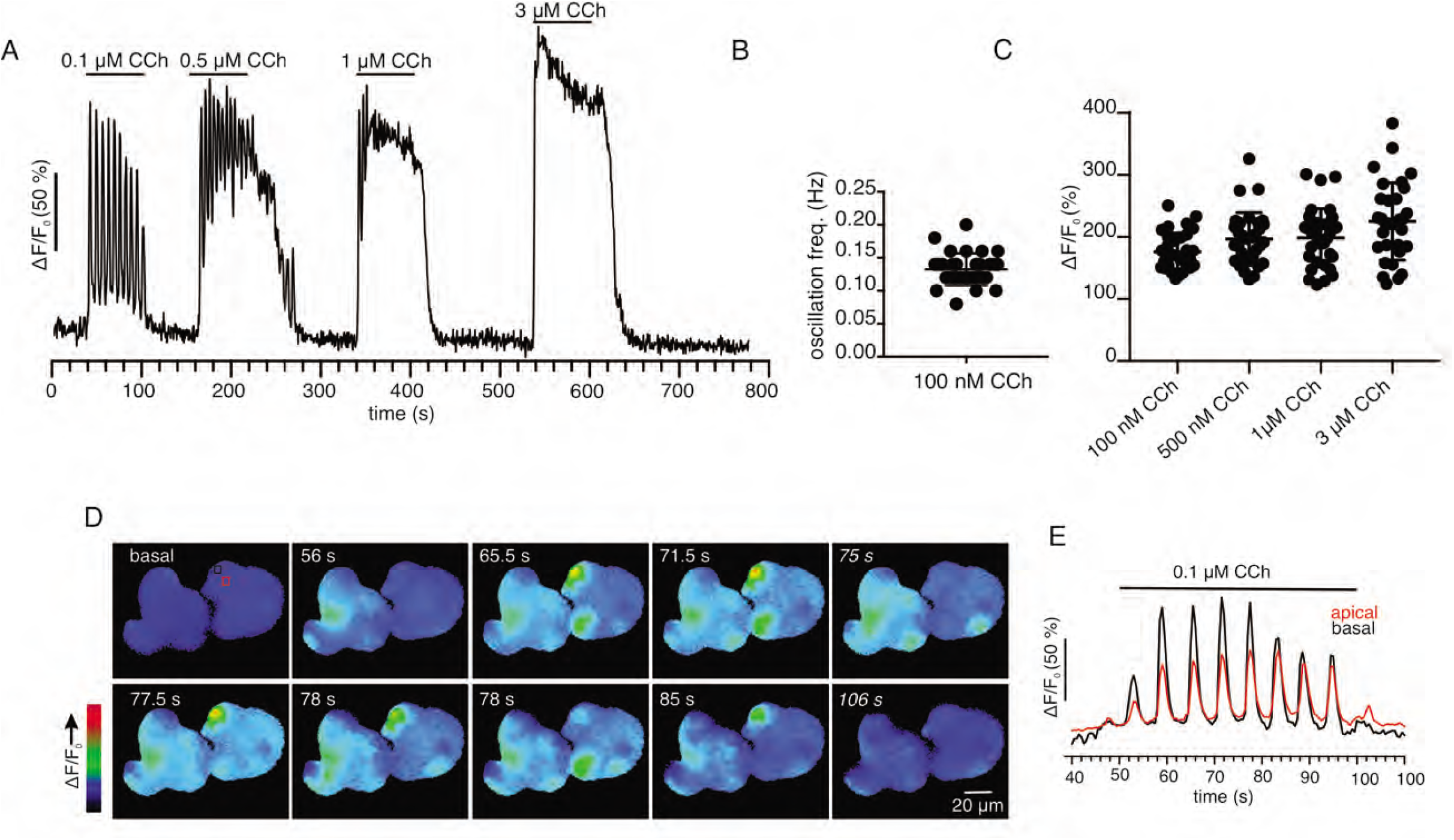
Ca^2+^ signaling in enzymatically isolated SMG acinar cells measured with GCaMP6f. **A**. A representative fluorescence changes evoked by superfusion of carbachol (CCh) from a single acinar cell within an acinus. **B**. Frequency of Ca^2+^ oscillations following exposure to 100 nM CCh. N = 32 cells from 3 animals. **C**. A plot of initial peak Ca^2+^ evoked by CCh. N = 32 cells from 3 animals. **D**. Time-course images of Ca^2+^ in response to CCh showing globalization of Ca^2+^ within a single cell. **E**. Ca^2+^ changes in the apical (red circle in the first frame of panel D) and basal (black circle in the first frame of panel D) domains of a single acinar cell.

Although GCaMP6f and fura-2 have similar affinities [7, 25], a possibility exists that the marked differences in the spatiotemporal properties of Ca^2+^ signals observed *in vivo* vs. documented *in vitro* are a function of the genetically encoded indicator used in the present studies. We therefore prepared small acinar clumps from Mist1^+/–^ x GCaMP6f^+/–^ animals by enzymatic digestion and monitored Ca^2+^ signals stimulated by the muscarinic agonist carbachol (CCh) by wide field imaging. At low concentrations of CCh, repetitive Ca^2+^ oscillations were evoked (Fig 10A) with a period of ∼0.15 Hz (Fig 10B, n=32 cells, 3 animals). In response to increasing concentrations of CCh, the initial peak tended to increase (Fig 10C, n=32 cells, 3 animals) and the oscillations dampened to generate an elevated plateau (Fig 10A). A statistically significant change in the initial peak height was not evident, possibly reflecting the relatively narrow CCh concentration range tested, or alternatively the relatively high affinity of GCaMP6f. Oscillations evoked by low concentrations of CCh were initiated in the apical region of the cell and invariably globalized (Fig 10D/E). In total, the spatiotemporal profile of the Ca^2+^ responses evoked in acutely isolated GCaMP6f expressing acini were strikingly similar to numerous reports documenting agonist-evoked Ca^2+^ responses with conventional chemical Ca^2+^ indicators and thus the properties of the genetically encoded indicator *per se* do not explain the differences observed.

We have previously constructed a series of mathematical models to help understand and explain our experimental results and those from other groups [1, 34, 42, 43, 50, 51]. Briefly, these multi scale models were based on cyclical global increases in [Ca^2+^] activating spatially separated ion channels required for sustained secretion in a 3D collection of seven acinar cells forming an acinus. Apical initiation of the Ca^2+^ signal results in activation of Cl^-^ flux through TMEM16a Cl channels and the Ca^2+^ signal is subsequently transmitted across the cell cytoplasm by a process of Ca^2+^-induced Ca^2+^ release (via RyR or IP3R). This leads to periodic increases of [Ca^2+^] in the basal aspects of the cell and hence activation of basal Ca^2+^ activated K^+^ (KCa) channels. Activation of the KCa channels is critical for maintenance of the electrochemical potential, driving the efflux of Cl^-^ and thus secretion. The spatiotemporal information from the current *in vivo* data is obviously not compatible with these models. We have therefore constructed a new model, as shown in Fig. 11. The apical region now harbors all the machinery necessary for saliva secretion, with KCa channels and Na/K ATPases in addition to the TMEM16a channels (Fig. 11) which are poised to respond to Ca^2+^ released predominately in the apical region, from IP3R that are situated in close proximity to the TMEM16A [34]. A more detailed description of the model can be found in methods and in [34, 50, 51].

**Fig. 11.**
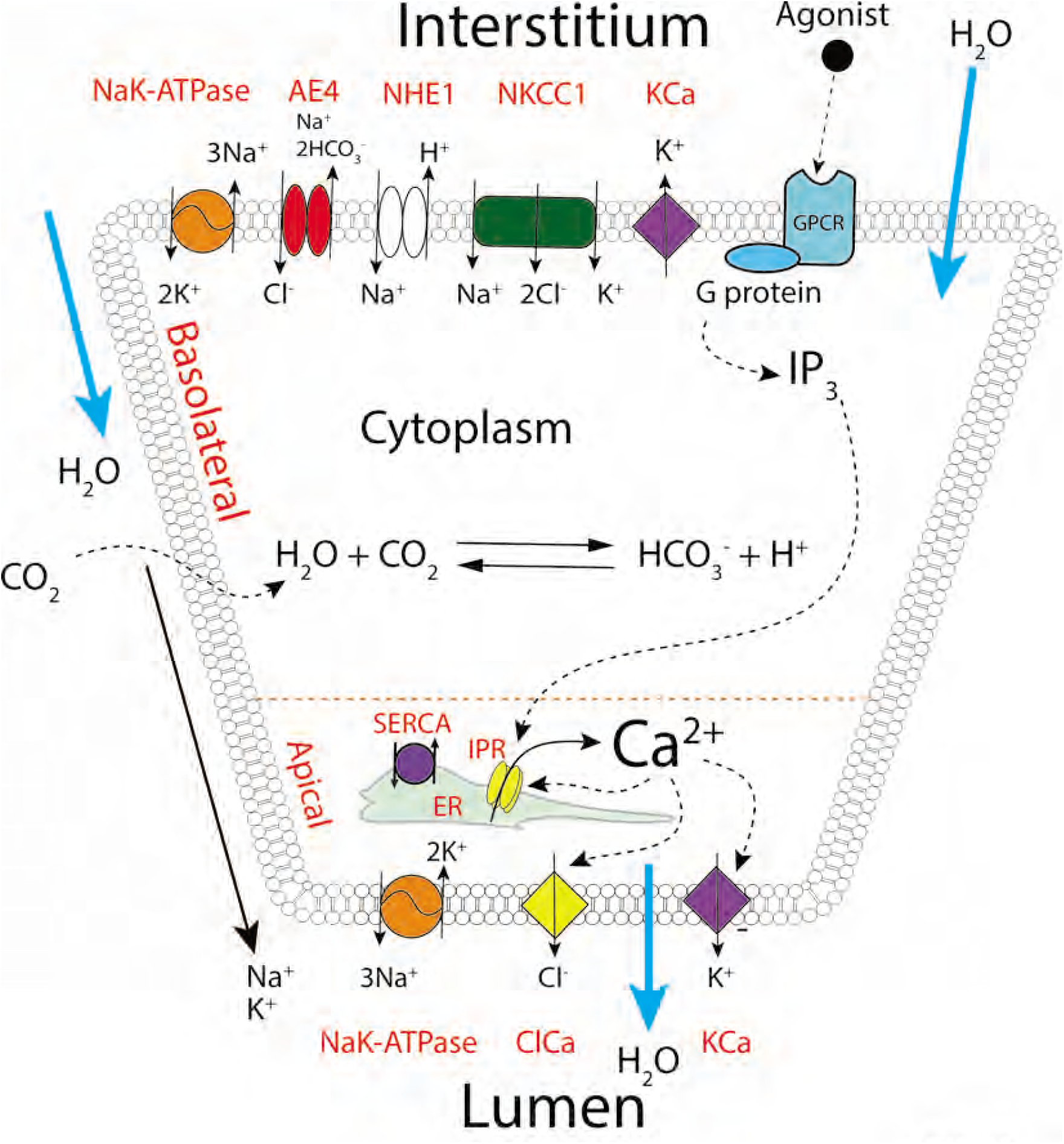
A proposed updated model for salivary secretion. The mathematical model is based on the fluxes and processes shown here. In contrast to previous models, the apical region now contains all the machinery needed for saliva secretion, included KCa channels and Na/K ATPases. Ca^2+^ is released predominately in the apical region, from IP_3_R that are situated in close proximity to TMEM16a, and there is no requirement for a propagated wave of increased [Ca^2+^] across the cell.

Results from the previous and current models are compared in Fig. 12. In the old model, agonist stimulation causes oscillatory fluid flow (panel A, green curve), resulting from apical [Ca^2+^] oscillations that are propagated across the cell, causing global periodic [Ca^2+^] oscillations (panel C). If the propagated [Ca^2+^] wave is removed, fluid secretion diminishes significantly (panel A, blue curve), as the [Ca^2+^] in the basal region exhibits little response (panel D). However, if KCa are now localized to the apical PM, together with Na/K-ATPases (to maintain a low [K^+^] in the primary saliva) then fluid section is restored (panel A, red curve) even though the [Ca^2+^] in the basal region is only marginally elevated (panel B).

**Fig 12.**
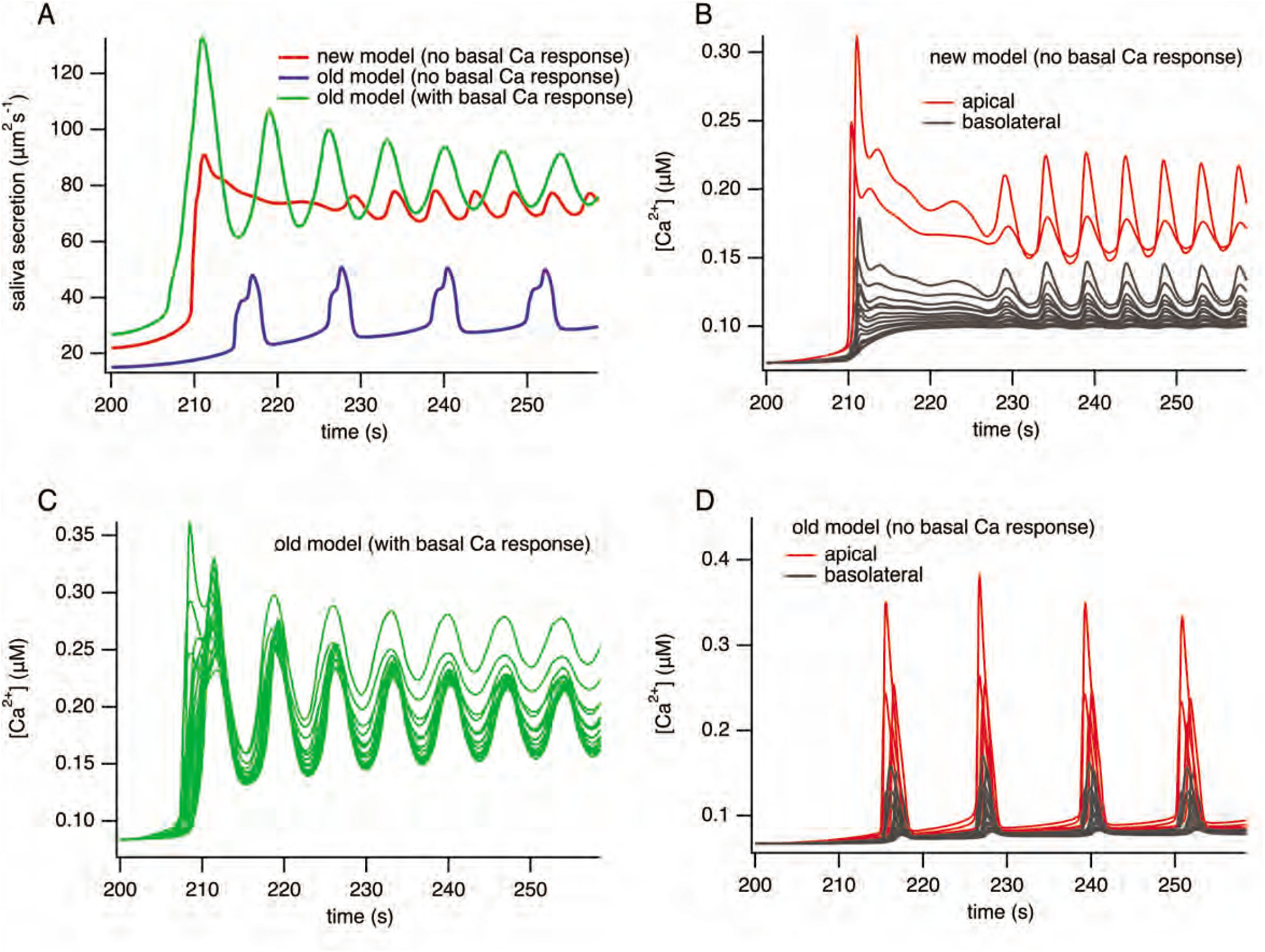
Comparison of results from the old and new models. The old model [35, 50, 51] has no KCa or NaK-ATPases in the apical membrane, but has a propagated wave of increased [Ca^2+^] across the cell. The new model has both KCa and NaK-ATPases in the apical membrane, but no propagated [Ca^2+^] wave. **A**, the old model with a propagated [Ca^2+^] wave gives normal fluid secretion (green curve) but if the propagated [Ca^2+^] wave is removed, fluid secretion is significantly diminished (blue curve) due to the lack of KCa activation. However, if the KCa are then put into the apical region (giving the new model, as shown by the red curve) fluid secretion is restored. **B**, [Ca^2+^] traces (from the new model) from randomly selected parts of the cell. The lower traces in grey are from the basal region, the upper traces in red are from the apical region. **C**. [Ca^2+^] traces (from the old model) from randomly selected parts of the cell. All the traces show raised [Ca^2+^], demonstrating the existence of a propagated [Ca^2+^] wave across the cell. **D**, [Ca^2+^] traces (from the old model) from randomly selected parts of the cell. In this version of the model the propagated [Ca^2+^] wave has been removed, as can be seen by the fact that the basal [Ca^2+^] responses in grey are much lower than the apical responses in red.

## Discussion

An increase in [Ca^2+^] following neurotransmitter release from parasympathetic neurons is fundamentally important for the stimulation of salivary fluid secretion. It is also recognized that the spatiotemporal properties of intracellular Ca^2+^ signals in exocrine cells are pivotal for appropriately activating ion channels necessary for the underlying process [2, 20, 21]. Despite a wealth of information documenting agonist-stimulated Ca^2+^ signals, recorded in dissociated single cells, isolated acini or excised lobules, the properties of physiological Ca^2+^ signals *in vivo* following endogenous neural stimulation have not been reported. We therefore generated transgenic mice that express a genetically encoded Ca^2+^ indicator specifically in exocrine acinar cells and used MP microscopy in live mice to document the Ca^2+^ signals generated following stimulation of the submandibular nerve which innervates the SMG. By establishing stimulation parameters optimum for fluid secretion, we have inferred the spatiotemporal properties of Ca^2+^ signals that ultimately underlie the physiological stimulation of fluid secretion at the lobule level and subsequently at subcellular resolution.

Following threshold stimulation, a small proportion of the imaged field responded after a prolonged latency by eliciting a single, or a few brief transients. These Ca^2+^ spikes were initiated in the extreme apical aspects of the cell, consistent with the localization of the majority of IP3R and previous *in vitro* data [24, 34, 54]. In stark contrast however, these signals did not globalize and were confined to the apical domain of individual single cells within an acinus. This threshold stimulation resulted in minor measurable fluid secretion. At greater stimulus strengths, where secretion was readily evident, an increasingly larger proportion of the field was recruited with shorter latency to yield multiple Ca^2+^ transients with increasing peak magnitude. The increase reflecting an increase in all cells within individual responding acini. Strikingly, these signals were again predominately localized to regions of the cell immediately juxtaposed to the apical PM with limited propagation towards the basal aspects of the cell. Upon optimal stimulation for secretion, essentially the entire field was recruited to produce a sustained increase with periodic fluctuations superimposed on the plateau after a minimal latent period. Notably, while the affinity of the indicator may have masked a greater magnitude of apical Ca^2+^ rise, or more prominent fluctuations in the signal at these higher stimulus strengths, a prominent apical to basal gradient was readily evident. Further increases in the intensity of the stimulation resulted in a rapid peak followed by a gradual waning of the signal and consistent with the absence of a sustained Ca^2+^ signal, the rate of fluid secretion was diminished. This reduced secretion likely results from a “tetanic” stimulus where either neurons fail to fire during their refractory period, neurotransmitter release is exhausted or acinar cell Ca^2+^ signaling is perturbed, during this likely non-physiological stimulation. Over the range of physiological stimulus strengths, the extent of fluid secretion was strongly correlated with the magnitude of the peak Ca^2+^ rise, rather than necessarily the frequency of Ca^2+^ transients. This conclusion is consistent with our previous modeling and experimental work that has indicated that the integrated increase in [Ca^2+^] during stimulation is the primary driving factor for efficient fluid secretion [42].

While local, non-propagating Ca^2+^ signals are reported in isolated pancreatic acinar cells, notably in whole-cell patch clamped cells where Ca^2+^ buffering is imposed [46, 47], the spatial characteristics of the Ca^2+^ signal observed *in vivo* in the present study represents a major difference to previous work in isolated salivary acinar cells. The failure of the signal to propagate and standing [Ca^2+^] gradients observed are presumably established by SERCA, PMCA and mitochondrial buffering to efficiently reduce the local [Ca^2+^] to levels that do not support substantial CICR through IP3R and RYR that are presumed to be localized in the ER distant from the apical trigger zone. An obvious question is, what factors or mechanisms underlie these differences? At the level of the SMG lobule, the polarity of individual cells and their organization within an acinus is vital for function. The complex architecture is maintained by junctional complexes and interactions with the extracellular matrix and stromal cells [8]. Clearly, these interactions may be disturbed when cells are isolated from an excised gland by enzymic digestion. An apparent consequence may be a disruption of the exquisite polarity of acinar cells. Perturbed localization of the Ca^2+^ signaling machinery could disrupt the balance between Ca^2+^ release and clearance to ultimately result in an alteration of the spatiotemporal characteristics of the signal. Conceivably, alterations in the polarization of the ER, such that IP3R or SERCA are not maintained in the extreme apical ER [23, 24], or a change in the distribution of mitochondria [11] to disrupt ER-mitochondrial junctions and effective Ca^2+^ sequestration from release sites could contribute to the altered Ca^2+^ signals observed *in vitro*.

A further important consideration when interpreting the characteristics of these Ca^2+^ signals is the obvious difference in delivery of the secretagogue in these experiments. In contrast to superfusion resulting in a defined “square” pulse of stimulating agent, as commonly used in *in vitro* studies, where in the acinar cells likely experience a complex, stimulus intensity, and time-dependent gradient of neurotransmitter. In our studies, the concentration of ACh released from neurons would be predicted to rapidly increase and then decay as it is hydrolyzed by acetylcholinesterases. As the frequency of stimulation is increased, the residual concentration of ACh present in the vicinity of acinar cells would be predicted to increase with a “saw-tooth” profile as the amount of ACh release exceeds the rate of degradation following more frequent, repetitive cycles of neurotransmitter exocytosis. In this scenario, when a threshold concentration of neurotransmitter is reached that generates sufficient IP3, an increase [Ca^2+^]i, will be triggered. The time to reach this threshold would define the latent period for that particular cell and the relative sensitivity to neurotransmitter will predicate whether an individual cell is responsive to a given stimulation. At optimum stimulation frequencies the residual ACh concentration would further increase and remain above the threshold for the majority of acinar cells to respond. This would facilitate greater Ca^2+^ release manifested as an increased peak and Ca^2+^ oscillations by the entire field. A summary of our observations and these ideas is presented in Supp Fig. 3. Notably, neural stimulation does not simply “pace” cells, as the frequency of oscillations does not correlate with the stimulus frequency. It is therefore likely, that the oscillations result from the inherent properties of IP3R and SERCA pumping activity but where the kinetics are influenced by the dynamic changes in neurotransmitter concentrations. This may contribute to the differences in oscillation frequency observed when comparing *in vivo* and *in vitro* data.

We observed considerable spatial heterogeneity in the responsiveness and peak amplitude of cells across the field of view. Specifically, there were sections of the field which exhibit the greatest sensitivity and largest responses at all stimulus intensities. Conceptually, this may reflect acini with expression of a higher density of cell surface receptors. Alternatively, highly responsive regions may represent cells physically closer to synapsing neurons and thus experiencing higher concentrations of ACh. The innervation of SMG lobules visualized in a reporter mouse expressing fluorescent protein in parasympathetic neurons has been reported to be fairly homogeneous [41]. Therefore, a further possibility is that spatial homogeneity arises in part because increasing numbers of axons are recruited to fire as stimulus intensity increases thus directly innervating a larger proportion of cells.

Clearly our previous models describing how Ca^2+^ signals promote salivary secretion are not consistent with the new *in vivo* data presented here. Fundamentally, if there is not an obligatory global intracellular Ca^2+^ wave, there is no means of activating the KCa channels in the basal PM that maintain salivary secretion. We have, however previously shown that the KCa channels, KCa3.1 (“IK”) and KCa1.1 (“BK”) together with Na/K ATPases are present in the apical membrane of salivary acinar cells [1, 2]. At the time these results were puzzling, as there was no obvious reason why the apical membrane should express either of these transport proteins. However, in combination with the new data presented here, the physiological importance of the apical KCa and Na/K ATPases is revealed. If [Ca^2+^] increases only in the apical region of the cell there must be KCa channels in the apical region, in order to maintain the electrochemical driving force for Cl^-^ secretion. On the other hand, since the primary saliva has a low [K^+^], this K^+^ must be removed from the lumen. This is the proposed function of the Na/K ATPase in the apical membrane.

Taken together, the present studies describe the spatial temporal characteristics of physiological Ca^2+^ signals driving salivary fluid secretion. Further, our findings highlight that caution should be exerted in extrapolating conclusions from *ex vivo* studies to physiological Ca^2+^ signals and function *in vivo*. Finally, it is envisioned that the present studies will provide a framework for investigating if Ca^2+^ signaling is disrupted in disease states such as Sjögren’s syndrome or radiation-induced xerostomia which result in pronounced hyposecretion of saliva [9, 45].

## Methods and Materials

Tamoxifen, corn oil, ketamine and Xylazine were purchased from Sigma Chemical, St Louis, Mo. Physostigmine was purchased from Tocris Minneapolis, Mn.

### Generation of mice expressing GCaMP6f in exocrine acinar cells

Adult female GCaMP6f^flox^ mice (Jackson Laboratory 029626) were crossed with male heterogenous Mist1^CreERT2^ (Jackson Laboratory 029228, kindly donated by Dr. Catherine Ovitt, University of Rochester) to generate Mist1^+/–^ x GCaMP6f^+/–^ transgenic mice. Tamoxifen (0.25 mg/g body weight) dissolved in corn oil was administered to mice at least 6 weeks old of either gender, by oral gavage once a day for three consecutive days. Imaging was carried out 1-4 weeks after the last tamoxifen administration. All animal procedures were approved by University Committee on Animal Resources (UCAR-2001-214E).

### *In vivo* MP imaging

The animal was anesthetized with Ketamine (80 mg/kg body weight, i.p.) and Xylazine (10 mg/kg body weight, i.p.). The animal was restrained with the ventral side up, and the animal’s body temperature was maintained at 37°C by a heat pad during surgery and imaging. A submandibular gland was exposed by making an incision in the skin on the ventral side of the neck. Connective tissue around the gland was teased away using forceps to allow the gland to be raised away from the body but remaining attached by a duct/nerve/blood vessels bundle. A pair of tungsten wires (WPI, Inc) was inserted to the bundle as stimulation electrodes. The lifted gland was placed on a 10 x 13 mm custom build small holder situated directly above the neck so that the position of the gland was not influenced by movement as a result of breathing. The gland was held by a cover glass to flatten the surface and to keep tissue moist with Hanks salt solution between the holder and the cover glass (Figure 1B). A 25x water immersion lens (Olympus XLPlan N 1.05 W MP) equipped with an objective heater (OKOLab COL2532) kept at 37°C was utilized for MP imaging. An upright 2-photon microscope system (FVMPE-RS, Olympus) with an excitation laser (InSight X3, Spectra-Physics) tuned at 950 nm and emission collected at 495-540 nm was utilized. Images were captured using Olympus FV31s-SW software at 30 fps framerate then averaged every 3 frames to achieve 10 fps image collection rate. Imaging depths between 10 and 55 µm from the surface of the gland were routinely utilized. PMT settings were fixed at 600V, 1x gain, and 3% black level, with excitation laser power adjusted per animal and according to imaging depth. Stimulation was generated by a stimulus isolator (Iso-flex, A.M.P.I.) set 5 mA, 200 μs, at the indicated frequency with train frequency and duration controlled by a train generator (DG2A, Warner Instruments). Images were captured typically for 30 s continuously. At least 1 min interval was given before imaging in the same field at a different stimulation strengths. In selected experiments, physostigmine (0.1 mg/kg body weight, i.p.) was administered to the animal on the microscope stage, at least 15 min prior to the imaging of post-physostigmine Ca^2+^ signals.

### Saliva secretion measurement

An anesthetized animal with stimulation electrodes attached to the duct bundle were gently held with ventral side up. A piece of filter paper, roughly 2 x 10 mm, were weighed before placing it in the animal’s mouth. A stimulation of 0–100 Hz was given to a gland for 1 min, and the paper was immediately removed and weighed. The difference of the weight of the paper represents the amount of saliva secreted out in the animal’s mouth. A new paper was placed in the mouth for each stimulation train.

### Acinar cell isolation and imaging

SMG acinar cells were enzymatically isolated from 2-4 month old, Mist1^+/–^ x GCaMP6f^+/–^ mice of both sexes. To isolate acinar cells, glands were extracted, connective tissue was removed, and glands were minced. Cells were placed in oxygenated dissociation media at 37°C for ∼30 minutes with shaking. Dissociation media consisted of Hank’s Balanced Salt Solution containing CaCl2 and MgCl2 (HBSS), bovine serum albumin (0.5%), and Collagenase Type II (0.2 mg/mL, Worthington). Cells were washed twice in HBSS with 0.5% BSA and resuspended in a HBSS solution containing 0.5% BSA and 0.02% trypsin inhibitor. Cells were then resuspended in imaging buffer (10 mM HEPES, 1.26 mM Ca^2+^, 137 mM NaCl, 4.7 mM KCl, 5.5 mM glucose, 1 mM Na_2_HPO_4_, 0.56 mM MgCl_2,_ at pH 7.4) and seeded onto a coverslip to allow attachment of cells. Cells were then perfused with imaging buffer and stimulated with agonist. Ca^2+^ imaging was performed using an inverted epifluorescence Nikon microscope with a 40 X oil immersion objective (NA=1.3). Cells were excited at 488 nm from a monochromator, and emission was monitored at 530 nm. Images were captured every 500 ms with an exposure of 10 ms and 4 × 4 binning using a digital camera driven by TILL Photonics software. Image acquisition was performed using TILLvisION software and data was exported to Microsoft excel.

### Batch Image analysis

Amplitude, latency, and percent of responding areas in fluorescence fields were analyzed using ImageJ software (NIH). XY drift during the imaging, when present, was corrected by applying the Descriptor-based series registration plugin (Stephan Preibisch). The imaging field was divided into 8 by 8 grids to yield 64 regions of interest (ROIs) of dimension 32×32 µm square. The average intensity of each ROI grid was generated for each frame. The average of first 100 frames prior to stimulation served as baseline fluorescence (F_0_), and %DF/F_0_ ((F– F_0_)/ F_0_ × 100) was calculated using the Image calculator function, so that the converted 32-bit image series represents [Ca^2+^] changes over time expressed as %DF/F_0_ in 8×8 alley of grids in the XY dimension (Fig. 5A). The standard deviation of the baseline 100 frames from grids in each image series provided an estimate of the noise level. A change greater than 4 times the standard deviation from the F_0_ value was considered as a stimulated Ca^2+^ signal and the ROI therefore designating a responding grid. To calculate latency prior to a response the time at the first incident Ca^2+^ signal meeting the criteria as a responding grid in each ROI after the initiation of the stimulation were recorded. Non-responding grids were excluded from this analysis. Statistical analyses were performed with paired t test, one-way ANOVA, and linear regression using Prism (GraphPad).

### Sub cellular image analysis

A computer based, automated region-of-interest (ROI) detection method that specifically targets cellular regions with pronounced changes was developed since no off-the-shelf tool met our particular requirements to efficiently process the data. The new software toolset consists of an assembly of Python (https://www.python.org) scripts deployed in a collection of Jupyter Lab (https://jupyter.org) “notebooks”. The notebooks were designed to automate repetitive analysis steps and give flexibility to interactively explore and analyze the image data sets and makes use of image processing algorithms in the scikit-image (https://scikit-image.org) package. ROI were identified by generating a difference image whereby the average image prior to stimulation was subtracted from the average image during the period of stimulation on a pixel-by-pixel basis (Figure 10A). An initial mask was created by simple pixel intensity thresholding (Figure 10B). Subsequent filtering by binary dilation and binary erosion removes undesired small regions and smooths the remaining regions (Figure 10C). The notebook then generates plots of pixel intensity over time for each ROI generated and utilizes the same mask for images sets generated from the same field at differing frequencies of stimulation (Figure 10D). A further script block was generated to objectively automate the identification of peaks generated by stimulation. For peak processing several signal processing operations from the scipy (https://www.scipy.org) package were employed. The script employs a sequence of signal resampling, low-pass zero phase filtering, high-pass zero phase filtering and a generic peak detector followed by a mapping back into the original response data. Detected peaks for each stimulation frequency were plotted in the notebook as black dots overlaid on region summary plots as shown in Figure 10D.

### Model details

The model has two interconnected modules; a Ca^2+^ oscillation module and a fluid secretion module. The two modules are coupled via changes in cell volume, and by the activation of KCa and ClCa by Ca^2+^. The Ca^2+^ oscillation module is a reaction-diffusion equation and is solved in a three-dimensional domain reconstructed from experimental structural data [51]. A finite element method was used. All other ions are assumed to be homogeneously distributed in the cell, and are thus described by a system of ordinary differential equations. The individual models for the various transporters, channels and exchangers are all given in [51]. The model equations were solved in a single, uncoupled cell (cell 4 in the notation of [51]). Similar results were obtained from all the other cells). The volume of the cell is taken into account, not by a full remeshing at each time step, but simply by a volume scaling factor in the reaction diffusion equation.

The Ca^2+^ oscillation module is based on a closed-cell Class I model in which oscillations arise via sequential activation and inactivation of the IP_3_R. [Ca^2+^] buffering is assumed to be fast and linear, and the IP_3_R model is taken from [43]. Ca^2+^ obeys a reaction-diffusion equation in the cell interior, but the effective diffusion coefficient of Ca^2+^ is small (due to buffering) which allows for localised increases in [Ca^2+^] which are not propagated throughout the cell. Because the IP_3_R are situated no more than 50 nm from the ClCa (which are on the apical membrane), release of Ca^2+^ through the IP_3_R is modelled as a boundary term, which avoids the necessity for a high-resolution finite element mesh in the apical region, which would greatly slow the calculations. The diffusion coefficient of IP_3_ is two orders of magnitude greater than that of Ca^2+^, and thus IP_3_ is effectively spatially homogeneous throughout the cell.

In the new model, all parameters and equations remain unchanged from those in the old model [51] with the following exceptions.

1. The maximum Ca^2+^ flux through RyR (V_RyR_) was set to 0. This is the simplest way of removing RyR from the model.
2. A K channel, with the same conductance as the basal KCa, was introduced into the apical membrane, with the equations for cytosolic and lumenal [K^+^] altered to compensate. The apical/basal K^+^ current ratio was determined by the apical/basal surface area ratio.
3. Na/KATPases were introduced into the apical membrane, using the same model and parameters as in (ref). The only change is that the total Na/KATPase activity was assumed to be unchanged, with 70% occuring in the basal membrane, and 30% in the apical membrane. The equations for cytosolic and lumenal [K^+^] and [Na^+^] were altered to compensate.
4. V_PLC,_ which controls the level of agonist stimulation, was set to 0.01 μM/s. As before [34, 50, 51], PLC activity was assumed to be dependent on [Ca^2+^] (although this has little effect on the oscillations), and nonzero only at the basal membrane.

## Supporting information

Manuscript

---

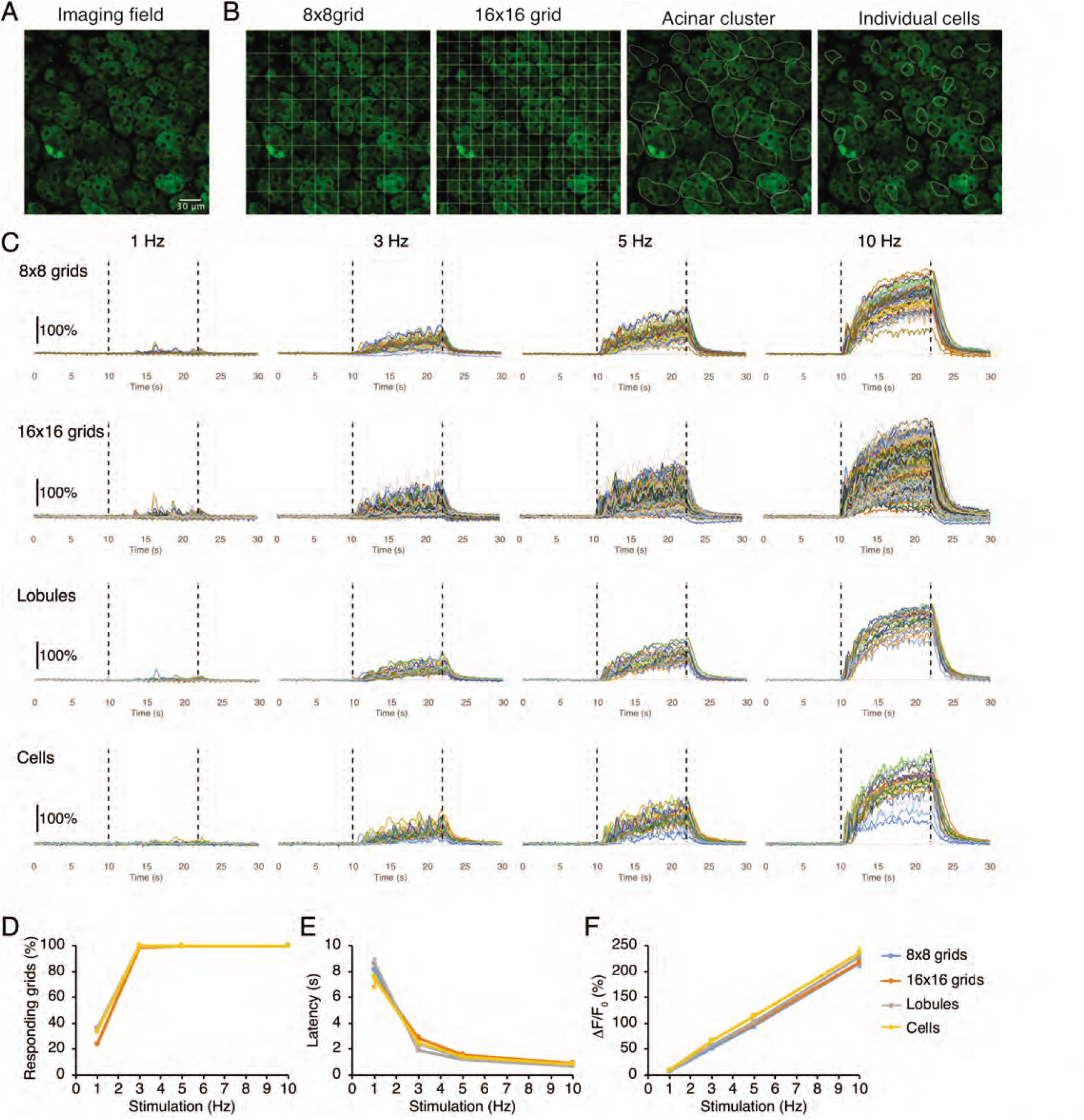

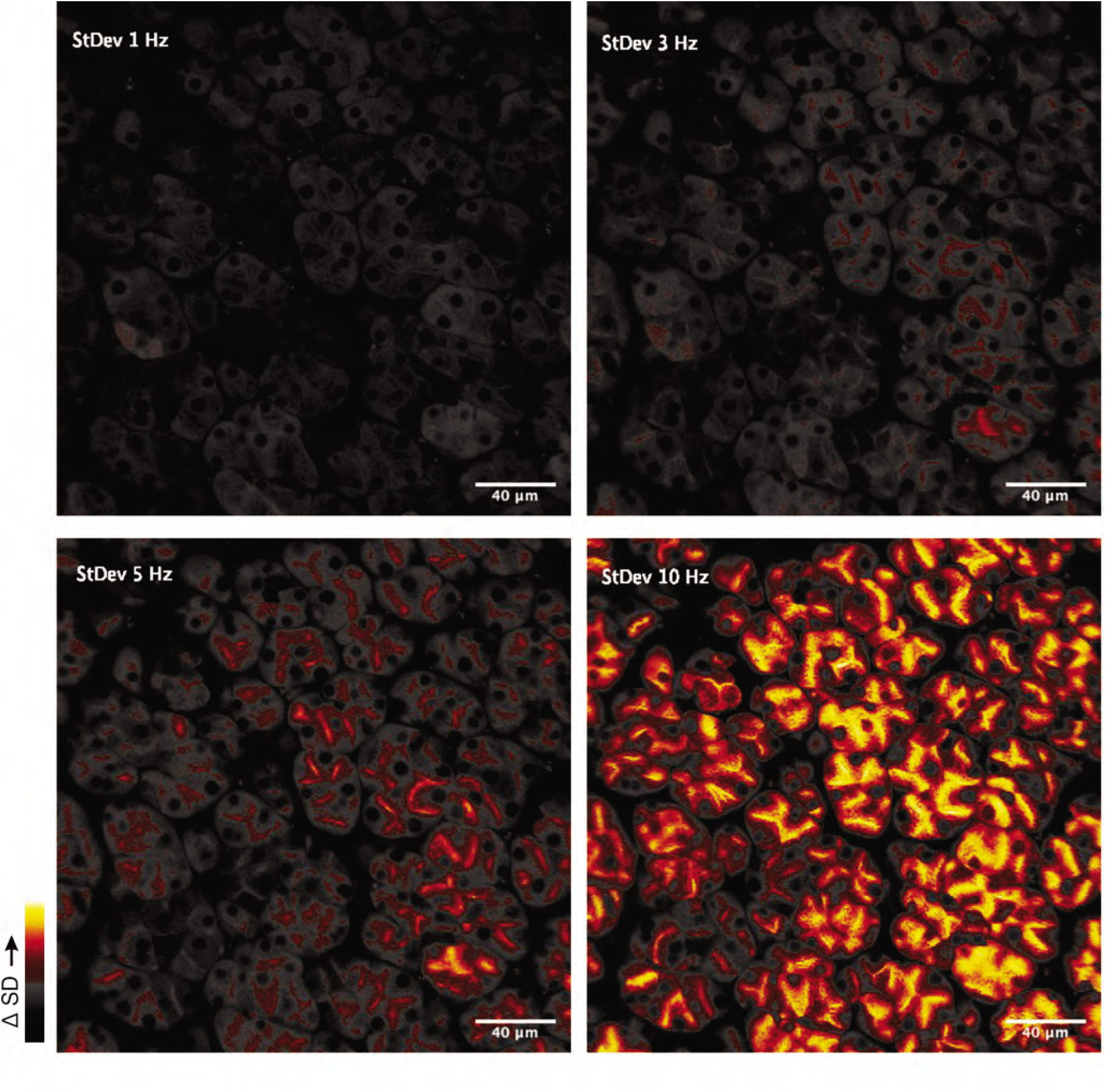

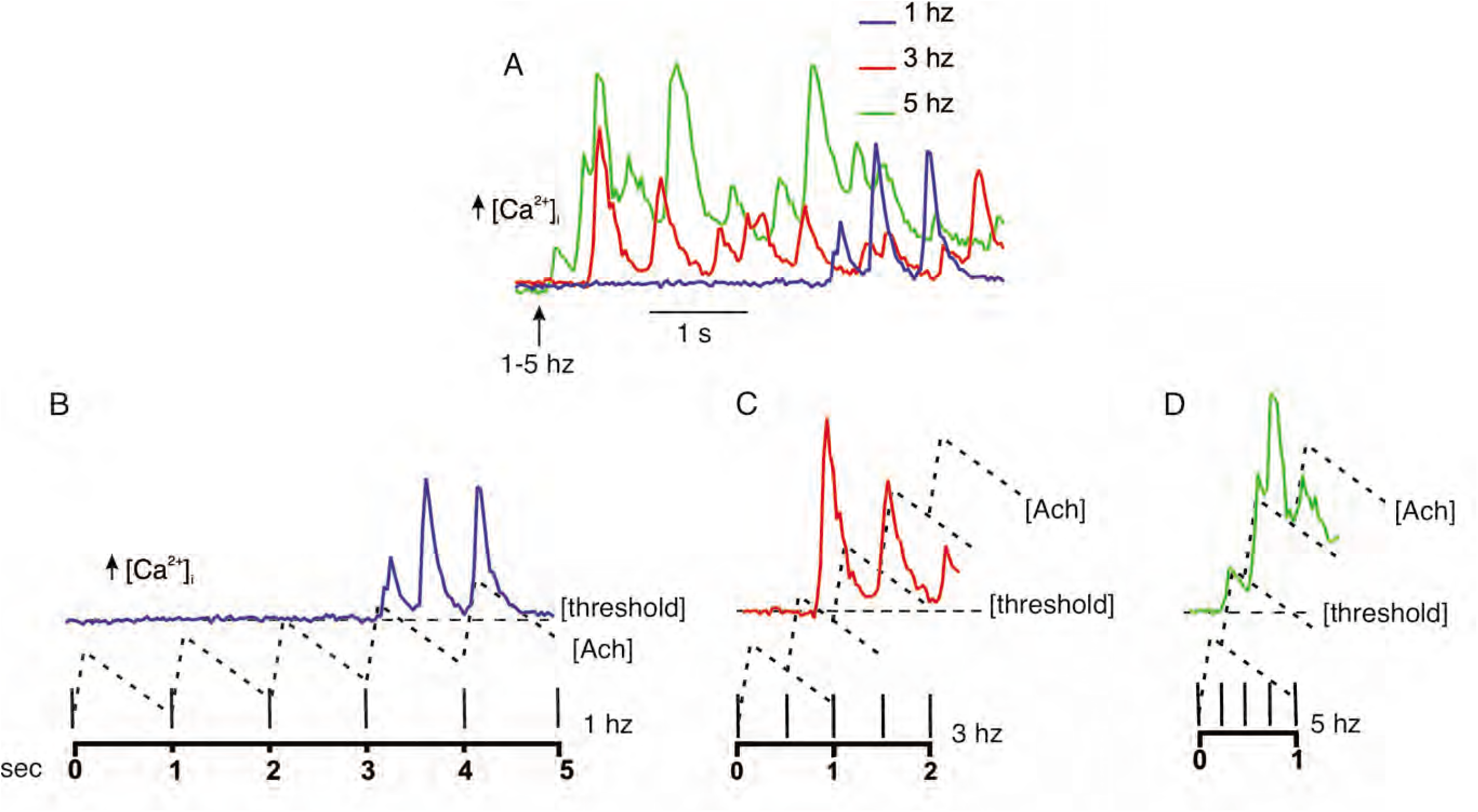

## References

1. Almassy, J, Siguenza, E, Skaliczki, M, Matesz, K, Sneyd, J, Yule, DI, and Nanasi, PP. New saliva secretion model based on the expression of Na(+)-K(+) pump and K(+) channels in the apical membrane of parotid acinar cells. Pflugers Arch 470(4): 613-621, 2018.

2. Almassy, J, Won, JH, Begenisich, TB, and Yule, DI. Apical Ca2+-activated potassium channels in mouse parotid acinar cells. J Gen Physiol 139(2): 121-133, 2012. PMC3269790

3. Ambudkar, IS. Dissection of calcium signaling events in exocrine secretion. Neurochem Res 36(7): 1212-1221, 2011. PMC3825030

4. Ambudkar, IS. Calcium signalling in salivary gland physiology and dysfunction. J Physiol 594(11): 2813-2824, 2016. PMC4887685

5. Arreola, J, Melvin, JE, and Begenisich, T. Activation of calcium-dependent chloride channels in rat parotid acinar cells. J Gen Physiol 108(1): 35-47, 1996. PMC2229297

6. Arreola, J, Park, K, Melvin, JE, and Begenisich, T. Three distinct chloride channels control anion movements in rat parotid acinar cells. J Physiol 490 (Pt 2): 351-362, 1996. PMC1158674

7. Badura, A, Sun, XR, Giovannucci, A, Lynch, LA, and Wang, SS. Fast calcium sensor proteins for monitoring neural activity. Neurophotonics 1(2): 025008, 2014. PMC4280659

8. Baker, OJ. Tight junctions in salivary epithelium. J Biomed Biotechnol 2010: 278948, 2010. PMC2825559

9. Baum, BJ, Alevizos, I, Zheng, C, Cotrim, AP, Liu, S, McCullagh, L, Goldsmith, CM, Burbelo, PD, Citrin, DE, Mitchell, JB, Nottingham, LK, Rudy, SF, Van Waes, C, Whatley, MA, Brahim, JS, Chiorini, JA, Danielides, S, Turner, RJ, Patronas, NJ, Chen, CC, Nikolov, NP, and Illei, GG. Early responses to adenoviral-mediated transfer of the aquaporin-1 cDNA for radiation-induced salivary hypofunction. Proc Natl Acad Sci U S A 109(47): 19403-19407, 2012. PMC3511089

10. Begenisich, T and Melvin, JE. Regulation of chloride channels in secretory epithelia. J Membr Biol 163(2): 77-85, 1998.

11. Bruce, JI, Giovannucci, DR, Blinder, G, Shuttleworth, TJ, and Yule, DI. Modulation of [Ca2+]i signaling dynamics and metabolism by perinuclear mitochondria in mouse parotid acinar cells. J Biol Chem 279(13): 12909-12917, 2004.

12. Bruce, JI, Shuttleworth, TJ, Giovannucci, DR, and Yule, DI. Phosphorylation of inositol 1,4,5-trisphosphate receptors in parotid acinar cells. A mechanism for the synergistic effects of cAMP on Ca2+ signaling. J Biol Chem 277(2): 1340-1348, 2002.

13. Chen, TW, Wardill, TJ, Sun, Y, Pulver, SR, Renninger, SL, Baohan, A, Schreiter, ER, Kerr, RA, Orger, MB, Jayaraman, V, Looger, LL, Svoboda, K, and Kim, DS. Ultrasensitive fluorescent proteins for imaging neuronal activity. Nature 499(7458): 295-300, 2013. PMC3777791

14. Cheng, KT, Ong, HL, Liu, X, and Ambudkar, IS. Contribution of TRPC1 and Orai1 to Ca(2+) entry activated by store depletion. Adv Exp Med Biol 704: 435-449, 2011. PMC3824974

15. Evans, RL, Park, K, Turner, RJ, Watson, GE, Nguyen, HV, Dennett, MR, Hand, AR, Flagella, M, Shull, GE, and Melvin, JE. Severe impairment of salivation in Na+/K+/2Cl-cotransporter (NKCC1)-deficient mice. J Biol Chem 275(35): 26720-26726, 2000.

16. Foskett, JK. [Ca2+]i modulation of Cl-content controls cell volume in single salivary acinar cells during fluid secretion. Am J Physiol 259(6 Pt 1): C998–1004, 1990.

17. Futatsugi, A, Nakamura, T, Yamada, MK, Ebisui, E, Nakamura, K, Uchida, K, Kitaguchi, T, Takahashi-Iwanaga, H, Noda, T, Aruga, J, and Mikoshiba, K. IP3 receptor types 2 and 3 mediate exocrine secretion underlying energy metabolism. Science 309(5744): 2232-2234, 2005.

18. Gallacher, DV and Petersen, OH. Stimulus-secretion coupling in mammalian salivary glands. Int Rev Physiol 28: 1-52, 1983.

19. Giovannucci, DR, Bruce, JI, Straub, SV, Arreola, J, Sneyd, J, Shuttleworth, TJ, and Yule, DI. Cytosolic Ca(2+) and Ca(2+)-activated Cl(-) current dynamics: insights from two functionally distinct mouse exocrine cells. J Physiol 540(Pt 2): 469-484, 2002. PMC2290247

20. Kasai, H and Augustine, GJ. Cytosolic Ca2+ gradients triggering unidirectional fluid secretion from exocrine pancreas. Nature 348(6303): 735-738, 1990.

21. Kidd, JF and Thorn, P. Intracellular Ca2+ and Cl-channel activation in secretory cells. Annu Rev Physiol 62: 493-513, 2000.

22. Larina, O and Thorn, P. Ca2+ dynamics in salivary acinar cells: distinct morphology of the acinar lumen underlies near-synchronous global Ca2+ responses. J Cell Sci 118(Pt 18): 4131-4139, 2005.

23. Lee, MG, Xu, X, Zeng, W, Diaz, J, Kuo, TH, Wuytack, F, Racymaekers, L, and Muallem, S. Polarized expression of Ca2+ pumps in pancreatic and salivary gland cells. Role in initiation and propagation of [Ca2+]i waves. J Biol Chem 272(25): 15771-15776, 1997.

24. Lee, MG, Xu, X, Zeng, W, Diaz, J, Wojcikiewicz, RJ, Kuo, TH, Wuytack, F, Racymaekers, L, and Muallem, S. Polarized expression of Ca2+ channels in pancreatic and salivary gland cells. Correlation with initiation and propagation of [Ca2+]i waves. J Biol Chem 272(25): 15765-15770, 1997.

25. Lewis, RS and Cahalan, MD. Mitogen-induced oscillations of cytosolic Ca2+ and transmembrane Ca2+ current in human leukemic T cells. Cell Regul 1(1): 99-112, 1989. PMC361429

26. Maruyama, EO, Aure, MH, Xie, X, Myal, Y, Gan, L, and Ovitt, CE. Cell-Specific Cre Strains For Genetic Manipulation in Salivary Glands. PLoS One 11(1): e0146711. 2016. PMC4709230

27. Maruyama, Y, Gallacher, DV, and Petersen, OH. Voltage and Ca2+-activated K+ channel in baso-lateral acinar cell membranes of mammalian salivary glands. Nature 302(5911): 827-829, 1983.

28. Matsui, M, Motomura, D, Karasawa, H, Fujikawa, T, Jiang, J, Komiya, Y, Takahashi, S, and Taketo, MM. Multiple functional defects in peripheral autonomic organs in mice lacking muscarinic acetylcholine receptor gene for the M3 subtype. Proc Natl Acad Sci U S A 97(17): 9579-9584, 2000. PMC16907

29. Melvin, JE. Saliva and dental diseases. Curr Opin Dent 1(6): 795-801, 1991.

30. Melvin, JE. Chloride channels and salivary gland function. Crit Rev Oral Biol Med 10(2): 199-209, 1999.

31. Melvin, JE, Yule, D, Shuttleworth, T, and Begenisich, T. Regulation of fluid and electrolyte secretion in salivary gland acinar cells. Annu Rev Physiol 67: 445-469, 2005.

32. Nakamoto, T, Romanenko, VG, Takahashi, A, Begenisich, T, and Melvin, JE. Apical maxi-K (KCa1.1) channels mediate K+ secretion by the mouse submandibular exocrine gland. Am J Physiol Cell Physiol 294(3): C810–819, 2008. PMC3298180

33. Nakamoto, T, Srivastava, A, Romanenko, VG, Ovitt, CE, Perez-Cornejo, P, Arreola, J, Begenisich, T, and Melvin, JE. Functional and molecular characterization of the fluid secretion mechanism in human parotid acinar cells. Am J Physiol Regul Integr Comp Physiol 292(6): R2380–2390, 2007.

34. Pages, N, Vera-Siguenza, E, Rugis, J, Kirk, V, Yule, DI, and Sneyd, J. A Model of [Formula: see text] Dynamics in an Accurate Reconstruction of Parotid Acinar Cells. Bull Math Biol 81(5): 1394-1426, 2019. PMC6449190

35. Pages, N, Vera-Siguenza, E, Rugis, J, Kirk, V, Yule, DI, and Sneyd, J. A model of calcium dynamics in an accurate reconstruction of parotid acinar cells. Bull Math Biol 81(5): 1394-1426, 2019. PMC6449190

36. Park, HS, Betzenhauser, MJ, Zhang, Y, and Yule, DI. Regulation of Ca(2)(+) release through inositol 1,4,5-trisphosphate receptors by adenine nucleotides in parotid acinar cells. Am J Physiol Gastrointest Liver Physiol 302(1): G97–G104, 2012. PMC3345966

37. Pedersen, AM, Bardow, A, Jensen, SB, and Nauntofte, B. Saliva and gastrointestinal functions of taste, mastication, swallowing and digestion. Oral Dis 8(3): 117-129, 2002.

38. Putney, JW, Jr. Inositol lipids and cell stimulation in mammalian salivary gland. Cell Calcium 3(4-5): 369-383, 1982.

39. Romanenko, VG, Catalan, MA, Brown, DA, Putzier, I, Hartzell, HC, Marmorstein, AD, Gonzalez-Begne, M, Rock, JR, Harfe, BD, and Melvin, JE. Tmem16A encodes the Ca2+-activated Cl-channel in mouse submandibular salivary gland acinar cells. J Biol Chem 285(17): 12990-13001, 2010. PMC2857126

40. Romanenko, VG, Nakamoto, T, Srivastava, A, Begenisich, T, and Melvin, JE. Regulation of membrane potential and fluid secretion by Ca2+-activated K+ channels in mouse submandibular glands. J Physiol 581(Pt 2): 801-817, 2007. PMC2075181

41. Sheu, SH, Tapia, JC, Tsuriel, S, and Lichtman, JW. Similar synapse elimination motifs at successive relays in the same efferent pathway during development in mice. Elife 6, 2017. PMC5315461

42. Sneyd, J, Crampin, E, and Yule, D. Multiscale modelling of saliva secretion. Math Biosci 257: 69-79, 2014. PMC4252247

43. Sneyd, J, Han, JM, Wang, L, Chen, J, Yang, X, Tanimura, A, Sanderson, MJ, Kirk, V, and Yule, DI. On the dynamical structure of calcium oscillations. Proc Natl Acad Sci U S A 114(7): 1456-1461, 2017. PMC5321031

44. Takemura, H and Putney, JW, Jr. Capacitative calcium entry in parotid acinar cells. Biochem J 258(2): 409-412, 1989. PMC1138377

45. Teos, LY, Zhang, Y, Cotrim, AP, Swaim, W, Won, JH, Ambrus, J, Shen, L, Bebris, L, Grisius, M, Jang, SI, Yule, DI, Ambudkar, IS, and Alevizos, I. IP3R deficit underlies loss of salivary fluid secretion in Sjogren’s Syndrome. Sci Rep 5: 13953, 2015. PMC4568516

46. Thorn, P, Lawrie, AM, Smith, PM, Gallacher, DV, and Petersen, OH. Local and global cytosolic Ca2+ oscillations in exocrine cells evoked by agonists and inositol trisphosphate. Cell 74(4): 661-668, 1993.

47. Thorn, P, Moreton, R, and Berridge, M. Multiple, coordinated Ca2+ -release events underlie the inositol trisphosphate-induced local Ca2+ spikes in mouse pancreatic acinar cells. EMBO J 15(5): 999-1003, 1996. PMC449994

48. Tojyo, Y, Tanimura, A, and Matsumoto, Y. Imaging of intracellular Ca2+ waves induced by muscarinic receptor stimulation in rat parotid acinar cells. Cell Calcium 22(6): 455-462, 1997.

49. Turner, RJ and Sugiya, H. Understanding salivary fluid and protein secretion. Oral Dis 8(1): 3-11, 2002.

50. Vera-Siguenza, E, Pages, N, Rugis, J, Yule, DI, and Sneyd, J. A Mathematical Model of Fluid Transport in an Accurate Reconstruction of Parotid Acinar Cells. Bull Math Biol 81(3): 699-721, 2019. PMC7219794

51. Vera-Siguenza, E, Pages, N, Rugis, J, Yule, DI, and Sneyd, J. A Multicellular Model of Primary Saliva Secretion in the Parotid Gland. Bull Math Biol 82(3): 38, 2020.

52. Warner, JD, Peters, CG, Saunders, R, Won, JH, Betzenhauser, MJ, Gunning, WT, 3rd, Yule, DI, and Giovannucci, DR. Visualizing form and function in organotypic slices of the adult mouse parotid gland. Am J Physiol Gastrointest Liver Physiol 295(3): G629–640, 2008. PMC2536791

53. Weiss, SJ, McKinney, JS, and Putney, JW, Jr. Receptor-mediated net breakdown of phosphatidylinositol 4,5-bisphosphate in parotid acinar cells. Biochem J 206(3): 555-560, 1982. PMC1158623

54. Won, JH, Cottrell, WJ, Foster, TH, and Yule, DI. Ca2+ release dynamics in parotid and pancreatic exocrine acinar cells evoked by spatially limited flash photolysis. Am J Physiol Gastrointest Liver Physiol 293(6): G1166–1177, 2007.

