## Supplementary material for "The characteristics of intracellular Ca^2+^ signals *in vivo* necessitate a new model for salivary fluid secretion": Manuscript

**SUPPLEMENTAL DATA.**

**Supplemental Fig. 1**. Grid subdivision of imaging field approximates the [$Ca^{2+}$] behaviors of lobules. A. A representative imaging field. B. 8x8 grids (1012 μ$m^2$ per grid) or 16x16 grids (253 μ$m^2$ per grid) were applied to the image. In addition, individual lobules (average = 1150 ± 65 μ$m^2$) and acinar cells (average 212 ± 12 μ$m^2$) were randomly chosen and manually selected as a reference. C. Time-course traces of [$Ca^{2+}$] in each grid/lobule before, during, and after 12s stimulations. D-F. Analysis results obtained from 8x8 grids and 16x16 grids agreed with the results from the lobules/cells.

**Supplemental Fig. 2.**

Standard Deviation images generated from the image series for the period of stimulation in a single represented imaged field**.** At all stimulus intensities the largest changes in fluorescence are observed in the extreme apical region of the acinar cluster. At lower stimulus intensities there is no appreciable propagation of the signal to the basal aspects of the cells. At higher intensities (10 Hz) a significant apical-basal gradient is established.

**Supplemental Fig. 3.** Proposed explanation for the temporal characteristics of $Ca^{2+}$ signals following neural stimulation. **A** shows the $Ca^{2+}$ changes following stimulation at the indicated intensities. With increasing intensity the latency is shortened and Ca2+ signals transition to a series of transients with increasing initial peak height. **B** the response to threshold stimulation. Neural stimulation results in the release of ACh which is degraded by acetylcholinesterases (doted line indicates [ACh]. When the [ACh] reaches a threshold sufficient to generate enough $IP_3$, a $Ca^{2+}$ signal is evoked. While the [ACh] is above this threshold further transients are evoked. **C-D** With higher frequency stimulation this threshold is reached quicker and the [ACh] is maintained to maintain $Ca^{2+}$ signals through the continued activity of $IP_3R$, refiling of ER pools and $Ca^{2+}$ clearance mechanisms.

**Supplemental Movie 1.** Movie generated by Python scripts running in the Jupyter lab environment following stimulation at 1 Hz. Left panel shows $\Delta F/F_0$ image series. Bottom right panel depicts the change in $F/F_0$ for the apical ROIs automatically generated by the software described in Figure 9.

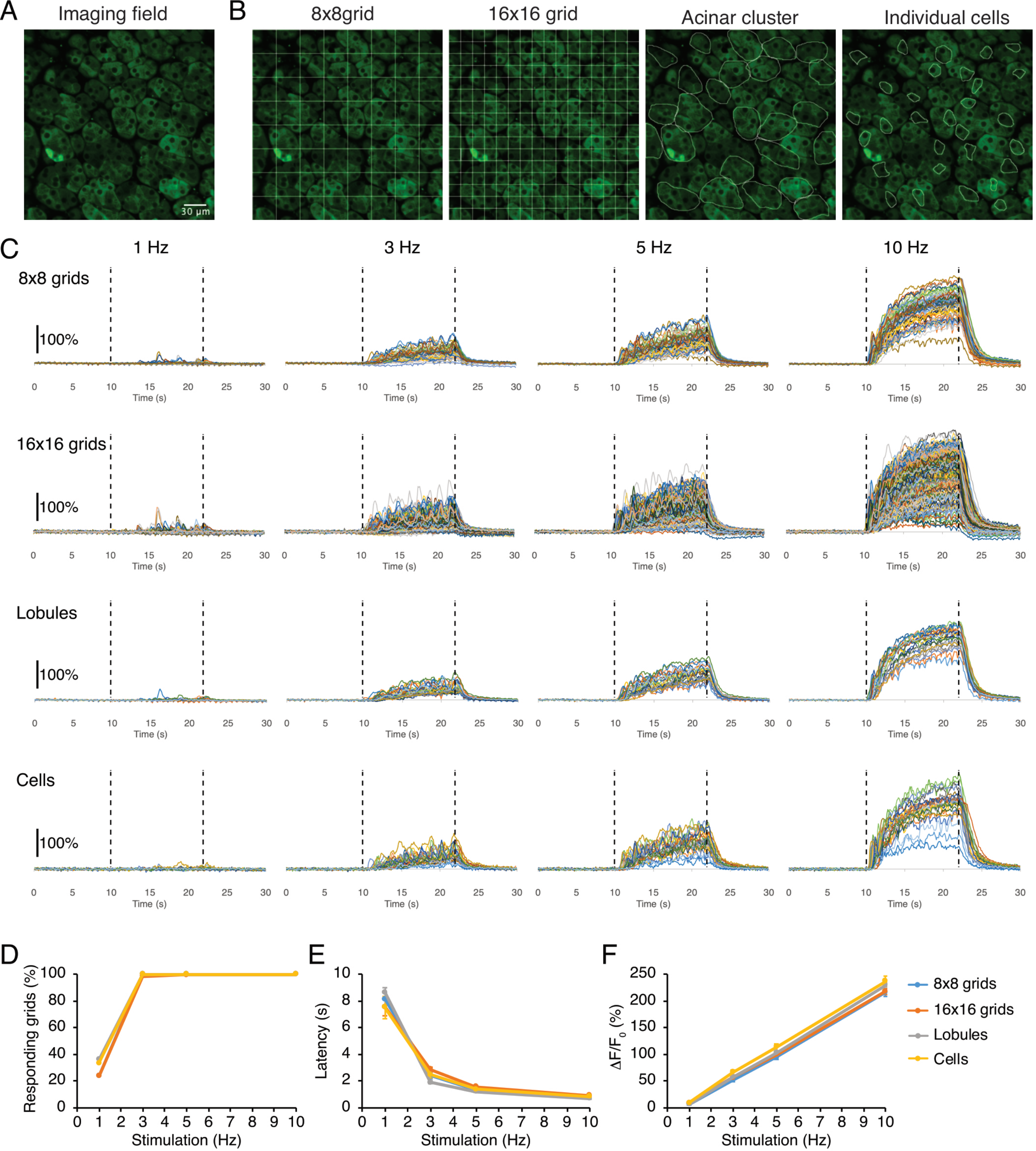
A
Imaging field
30 µm
B
8x8grid
16x16 grid
Acinar cluster
Individual cells
C
1 Hz
3 Hz
5 Hz
10 Hz
8x8 grids
16x16 grids
Lobules
Cells
100%
Time (s)
D
Responding grids (%)
Stimulation (Hz)
E
Latency (s)
Stimulation (Hz)
F
ΔF/F0 (%)
Stimulation (Hz)
8x8 grids
16x16 grids
Lobules
Cells

supp 1

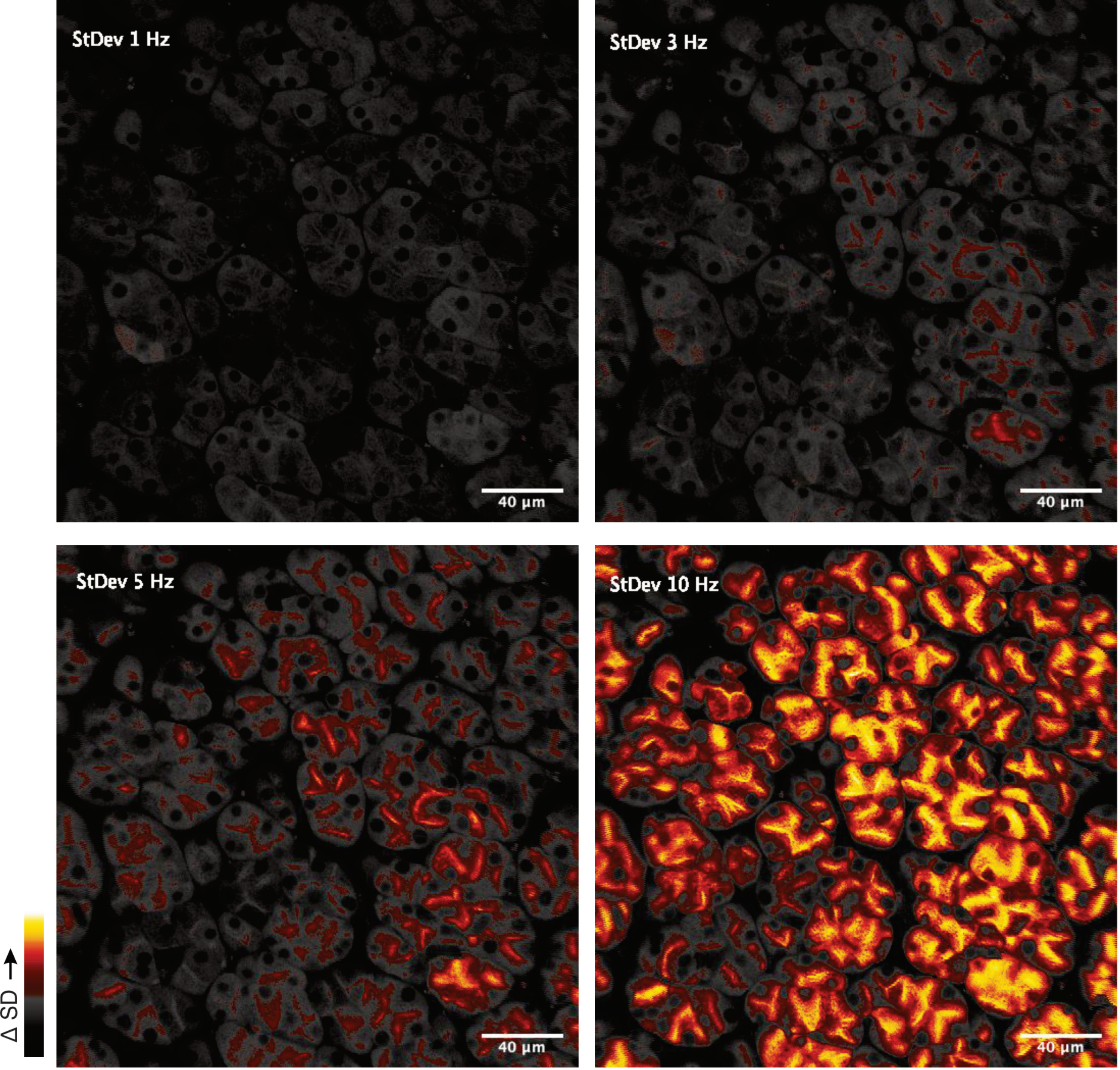
StDev 1 Hz
40 µm
StDev 3 Hz
40 µm
StDev 5 Hz
40 µm
StDev 10 Hz
40 µm
Δ SD

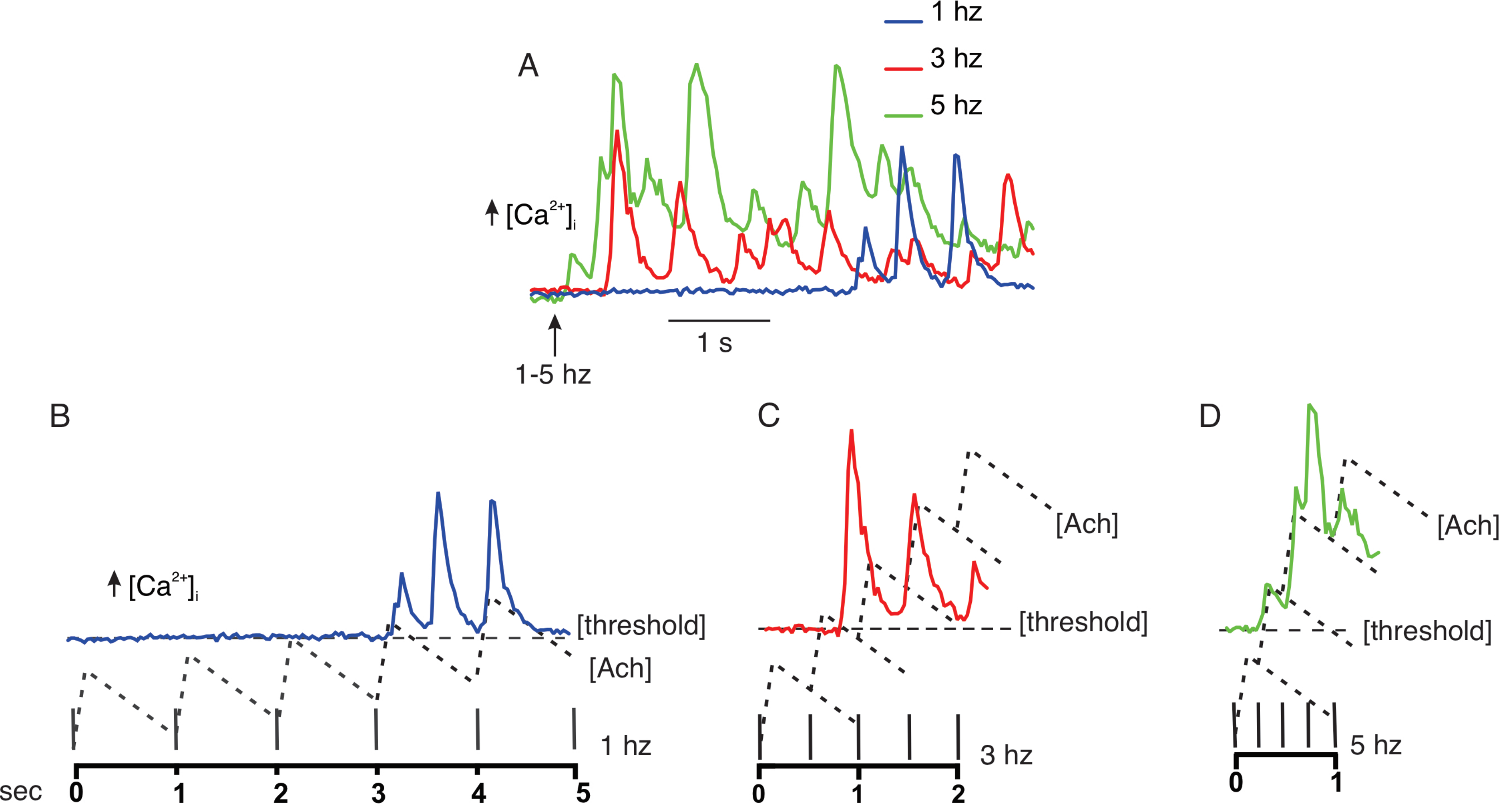
A
1 hz
3 hz
5 hz
[Ca2+]i
1 s
1-5 hz
B
[Ca2+]i
[threshold]
[Ach]
1 hz
sec
0
1
2
3
4
5
C
[Ach]
[threshold]
3 hz
0
1
2
D
[Ach]
[threshold]
5 hz
0
1

SUPP3
